# ER stress and cell fate: FKBP2 regulates proinsulin folding and α- vs. β cell differentiation via NFAT and HDAC9

**DOI:** 10.1101/2025.09.01.672695

**Authors:** Edyta Urbaniak, Maja Baginska, Wojciech J. Szlachcic, Malgorzata Grabowska, Ekaterina Shcheglova, Artur Jankowski, Michal T. Marzec, Malgorzata Borowiak

## Abstract

The continuous demand for insulin production places a burden on the endoplasmic reticulum (ER) of pancreatic β cells, making them highly susceptible to protein misfolding and ER stress. While general ER chaperones are known to be essential for β cell homeostasis, the specific mechanisms by which individual chaperones facilitate proinsulin folding and influence cell fate remain unclear. This study identifies FK506-binding protein 2 (FKBP2), an ER-localized cis-trans prolyl isomerase, as a critical regulator of human β cell differentiation and proinsulin processing. Using loss- and gain-of-function FKBP2 models in human pluripotent stem cells (hPSCs), we demonstrate that its deficiency impairs insulin processing, leading to abnormal insulin granule morphology and reduced insulin secretion. Unexpectedly, our findings reveal a crucial and unreported role for FKBP2 in cell fate determination. Single-cell RNA sequencing reveals that FKBP2 loss disrupts the proper endocrine lineage allocation, causing a shift towards the α cell formation at the expense of β cells. In KO endocrine cells, we observed sustained ER stress and elevated intracellular calcium levels, along with activation of the NFAT2–HDAC9 axis. Pharmacological inhibition of HDAC class IIa activity partially rescued β cell differentiation, supporting a causal role for this pathway. Collectively, our results provide new mechanistic insights into how ER chaperones can control pancreatic development and contribute to the pathogenesis of diabetes.

## Introduction

Precise regulation of blood glucose homeostasis depends upon efficient insulin biosynthesis. Insulin, synthesized by pancreatic β cells in the islets of Langerhans ^1^, plays a pivotal role in preventing hyperglycemia and attenuating long-term diabetic complications. Comprising only an estimated 1 g of total weight in humans, pancreatic β cells are a vulnerable link in metabolism. The β cells exhibit a high biosynthetic capacity, synthesizing up to 6,000 proinsulin molecules per second. The biosynthetic pathway commences with the synthesis of preproinsulin, a single-stranded precursor molecule. Subsequent signal peptide cleavage upon entering the endoplasmic reticulum (ER) results in the formation of proinsulin, a polypeptide comprising α, β, and C-peptide domains. Within the rough ER, proinsulin folds, and three intramolecular disulfide bonds are created between the α and β chains, a process facilitated by protein disulfide isomerase ^2–4^. Subsequently, proinsulin dimerizes and is transported to the Golgi apparatus, where it undergoes further assembly, a process facilitated by zinc, calcium, and an acidic environment. In secretory granules, proinsulin hexamers are cleaved by the endoproteases prohormone convertase 1/3 and 2 (PC1/3 and 2), to yield and store mature insulin. Upon an increase in blood glucose levels, these granules release insulin into the bloodstream via exocytosis.

Proinsulin production constitutes a significant load on β cell ER, and decompensated endoplasmic reticulum (ER) stress is a cause of β cell failure and loss in both type 1 diabetes (T1D) and type 2 diabetes (T2D). Even under physiological conditions, approximately 30% of nascent insulin may exhibit misfolding, resulting in compromised functionality ^5,6^. The intracellular accumulation of misfolded insulin within the ER impedes secretory capacity, thereby contributing to the development of hyperglycemia. This hyperglycemic state imposes an augmented biosynthetic demand upon β cells, necessitating increased proinsulin synthesis. Higher proinsulin synthesis exacerbates protein misfolding and induces ER stress, establishing a detrimental positive feedback loop. This cyclical process ultimately culminates in β cell dysfunction and apoptotic cell death, leading to the progression of diabetes ^7^. Despite ongoing research, the precise chaperone-mediated mechanisms governing proinsulin folding within β cells remain incompletely understood.

Interestingly, the protein folding mechanism exhibits evolutionary conservation across the vertebrate insulin and insulin-like growth factor (IGF) superfamily, where proinsulin and IGF-2 share both conserved and divergent structural characteristics, folding prerequisites, and overlapping functionalities. By elucidating this pathway, we identified FK506-binding protein 2 (FKBP2), also known as FK506-binding protein 13, an ER chaperone and *cis-trans* proline isomerase, as a proinsulin-specific factor that has evolved to constitute the high-throughput proinsulin folding system ^8^. As a member of the peptidyl-prolyl cis/trans isomerase (PPIase) superfamily, FKBP2 binds to proinsulin and catalyzes X-Pro peptide bond isomerization. This catalytic activity is essential for proper protein folding, as proline isomerization critically alters protein conformation, thereby influencing its activity and ligand recognition. FKBP2 expression is elevated in response to the accumulation of unfolded proteins during ER stress in transformed rat and human β cell lines ^8^. Compellingly, FKBP2 has been implicated in T2D through proteomic analyses, and genetic polymorphisms in FKBP2 are associated with an increased susceptibility to T2D.

Here, we investigate the role of FKBP2 in human β cells using human pluripotent stem cells (hPSCs) pancreatic differentiation. We observed that FKBP2 is expressed in endocrine cells both *in vitro* and *in vivo*, which prompted us to investigate the role of FKBP2 in proinsulin folding. Unexpectedly, our investigation revealed the FKBP2 expression during β cell development, specifically within newly formed β cells and endocrine progenitors (EP). This finding suggests a potentially novel and crucial role for FKBP2 in β cell differentiation. Interestingly, another chaperone, GPR94, a HSP90-like protein localized in the ER, known for its specialized function in protein folding and quality control, was shown to regulate β cell development, possibly through its interaction with and control over the folding of IGF1/2 or integrin proteins ^8^. Here we demonstrate that FKBP2 is not only essential for proinsulin folding in human β cells but also represents a critical factor in the complex molecular network orchestrating β cell development.

## Results

### FKBP2 expression in human pancreas and hPSC-derived β cells

We have previously reported the expression of FKBP2 in human β cells *in vivo* ^8^. Building on this, we investigated the FKBP2 expression in the human adult pancreas. Immunofluorescence staining revealed FKBP2 presence in the majority of insulin-positive β cells. In contrast, somatostatin-positive (SST+) δ cells rarely co-expressed with FKBP2 (**Fig. 1A**). Interestingly, some glucagon-positive (GCG+) α cells also exhibited FKBP2 immunoreactivity (**Fig. S1A**). We also detected *FKBP2* mRNA expression in endocrine cells, including β cells in human adult pancreas single-cell RNA-sequencing (scRNA-seq) datasets ^9^ (**Fig. S1B**).

**Figure 1:**
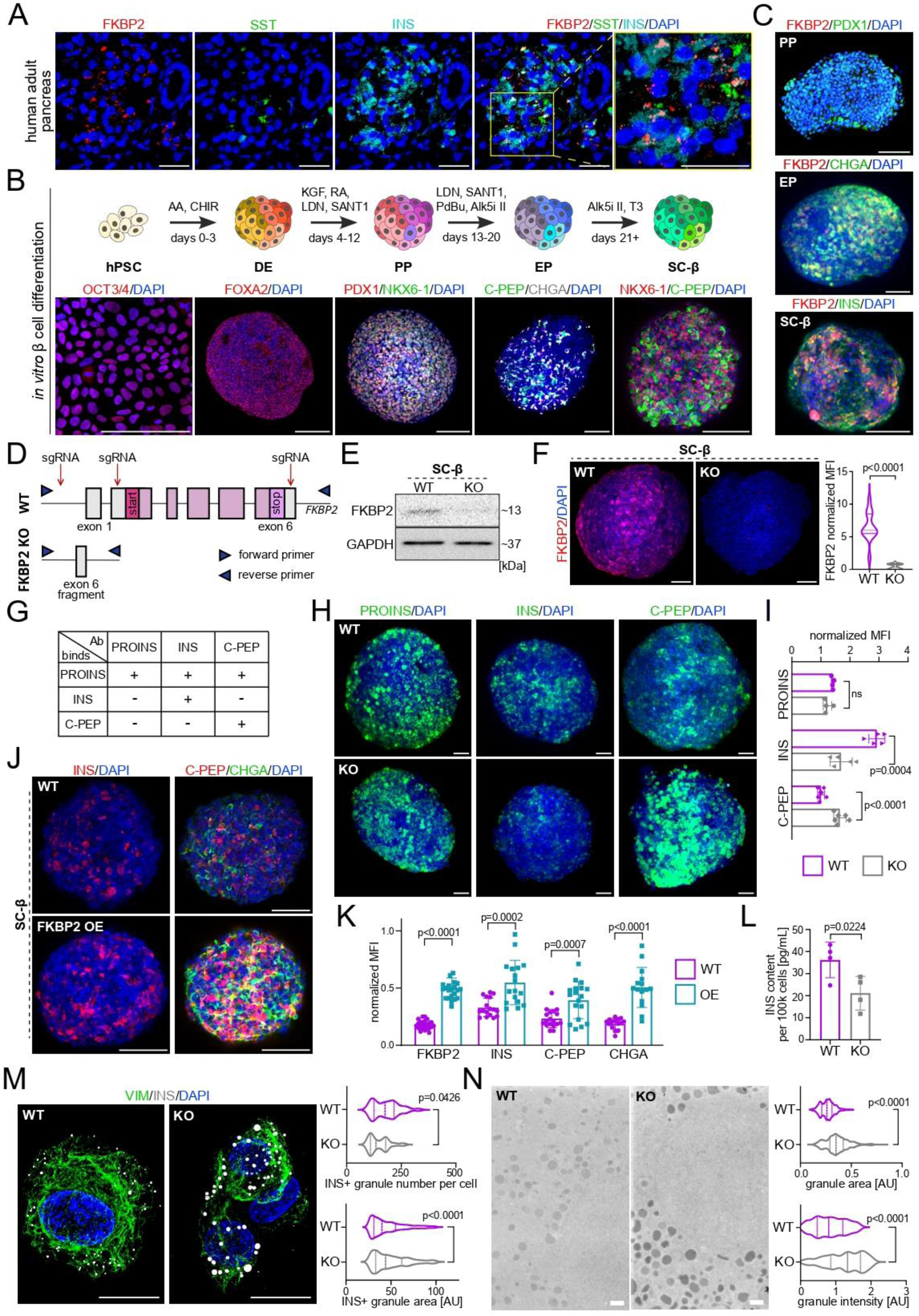
FKBP2 is expressed in pancreatic β cells, and its deletion impacts insulin processing and granule morphology in human SC-β cells. **(A)** Immunofluorescence staining of the human adult pancreas sections for FKBP2 (red), somatostatin (SST, green), and insulin (INS, cyan). DAPI marks nuclei (blue). The boxed region indicates an area of interest with co-localization, and a magnified view is shown on the far right. Scale bar, 50 µm. **(B)** Schematic of the *in vitro* differentiation protocol of human pluripotent stem cells (hPSCs) into pancreatic β cells (SC-β cells). Representative immunofluorescence images show expression of key markers at different stages: OCT3/4 (red, hPSCs), FOXA2 (red, definitive endoderm, DE), PDX1 and NKX6-1 (red and green, pancreatic progenitors, PP), CHGA (white, endocrine progenitors, EP), as well as NKX6-1, INS, and C-PEP (SC-β cells, SC-β). Scale bar, 100 µm. **(C)** Immunofluorescence staining of FKBP2 during *in vitro* β cell differentiation. FKBP2 (red) expression is detected in PPs (PDX1+, green), EPs (CHGA+, green), and SC-β cells (INS+, green). DAPI marks nuclei (blue). Scale bar, 100 µm. **(D)** Diagram illustrating CRISPR/Cas9-mediated FKBP2 knockout strategy in hPSCs. The entire FKBP2 gene was targeted for deletion. sgRNA target sites in exons 1, 2 and 6 of the FKBP2 gene, along with the locations of PCR primers used for validation, are indicated. **(E)** Western blot analysis of FKBP2 protein expression in wild-type (WT) and FKBP2 KO SC-β cells. GAPDH serves as a loading control. **(F)** Representative immunofluorescence staining of FKBP2 (red) expression in WT and FKBP2 KO SC-β cells. DAPI marks nuclei (blue). The violin plot on the right shows the quantification of the normalized mean fluorescence intensity (MFI) of FKBP2 across replicates. The plot displays the data distribution, with the median (dashed line) and interquartile range (IQR, dotted line, representing the 25th–75th percentiles). N = 3 biological replicates. The p-value was determined using an unpaired t-test. Scale bar, 100 µm. **(G)** The table presents the antibody binding specificities, showing binding patterns of proinsulin (PROINS), insulin (INS), and C-peptide (C-PEP) antibodies. Positive (+) and negative (-) reactivity indicated. **(H)** Immunofluorescence staining of proinsulin (PROINS, green, left), insulin (INS, green, center), and C-peptide (C-PEP, green, right) in WT and KO SC-β cells. DAPI marks nuclei (blue). Scale bar, 100 µm. **(I)** Quantification of normalized mean fluorescence intensity (MFI) of proinsulin (PROINS), insulin (INS), and C-peptide (C-PEP) in WT and FKBP2 KO SC-β cells. Each dot represents one image; horizontal lines indicate mean ± SD. N = 3 biological replicates, an unpaired t-test was used to assess significance, and p-values are indicated. **(J)** Immunofluorescence staining for insulin (INS, red), chromogranin A (CHGA, green), and C-peptide (C-PEP, red) in WT and FKBP2 overexpressing (OE) SC-β cells. DAPI marks nuclei in blue. FKBP2 OE was induced by 1 µg/mL doxycycline treatment at day 20 of differentiation and cells are analyzed at day 26. Scale bar, 100 µm. **(K)** Quantification of FKBP2, insulin (INS), C-peptide (C-PEP), and chromogranin A (CHGA) protein expression levels in WT and FKBP2 overexpressing (OE) SC-β cells at day 26 of differentiation. The OE hPSC line was treated with 1 µg/mL doxycycline on day 20 of differentiation. Each dot represents one image (4-6 images per biological repeat, N = 3 independent biological repeats), with mean and SD indicated. An unpaired t-test was used to determine statistical significance. **(L)** Total insulin secretion following stimulation with 30 mM KCl per 100k WT and FKBP2 KO SC-β cells, at day 46. Each dot represents one biological replicate run in two technical replicates, with mean and SD indicated, N = 3 independent biological replicates. An unpaired t-test was used to determine statistical significance. **(M)** Immunofluorescence staining and SIM images of WT and FKBP2 KO SC-β cells show expression of vimentin (VIM, green) and insulin (INS, white). DAPI marks nuclei (blue). Scale bar, 10 µm. Quantifications of INS granule number and area are shown on the right, as violin plots. The median is indicated as a dashed line and IQR as a dotted line, representing 25th–75th percentiles. Statistical significance was assessed using unpaired t-test; p-values are indicated on the plot, N = 3 independent biological replicates. **(N)** Representative transmission electron microscopy micrographs of WT and FKBP2 KO SC-β cells display granules in WT and KO cells. Quantifications of insulin granule size and counterstaining signal intensity (AU, arbitrary units) are shown as violin plots. The violin plots show the data distribution with the median (dashed line) and IQR (dotted line, representing 25th– 75th percentiles). N = 3 independent biological replicates. An unpaired t-test was used to determine statistical significance. Scale bar, 2 µm.

To understand FKBP2 role in human β cells, we utilized directed pancreatic differentiation of hPSCs using a previously established protocol with modifications ^10–12^. During the *in vitro* differentiation cells progressively adopt the fate of definitive endoderm (DE), pancreatic progenitors (PP), EP, and finally, β cells (SC-β) (**Fig. 1B**). Immunostaining and flow cytometry for stage-specific markers, including POU class 5 homeobox 1 (OCT3/4) for hPSCs, forkhead box A2 (FOXA2) for DE, pancreatic and duodenal homeobox 1 (PDX1) and NK6 homeobox 1 (NKX6-1) for PPs, chromogranin A (CHGA) for EPs, and C-peptide (C-PEP), insulin and NKX6-1 for SC-β cells confirmed the successful various pancreatic progenitors and SC-β cells generation (**Fig. 1B** and **S1C**). Analysis of publicly available scRNA-seq datasets of *in vitro* differentiating β cells revealed *FKBP2* mRNA expression in EPs and SC-β cells ^13^ (**Fig. S1D**). Using immunofluorescence staining and western blot, we detected FKBP2 protein starting from the EP stage, and FKBP2 expression was maintained in CHGA⁺ and INS⁺ early (day 21) and late (day 31) SC-β cells (**Fig. 1C** and **S1E-F**). Together, these results suggest that FKBP2 is present when endocrine β cells are formed *in vitro*.

### Impact of FKBP2 loss- and gain-of function on insulin processing and secretion

To investigate the role of FKBP2 in SC-β cells, we generated FKBP2 knockout (KO) hPSCs using CRISPR-Cas9 gene editing (**Fig. 1D**). The deletion of the entire FKBP2 gene was confirmed at the DNA level by PCR (**Fig. S1G)** and Sanger sequencing (**Fig. S1H**). Additionally, we verified the absence of FKBP2 at the protein level in SC-β cells using western blot (**Fig. 1E** and **S1I**) and immunofluorescence staining (**Fig. 1F)**. We observed no major effect on the growth of KO hPSCs, as monitored by live-cell imaging during a 5-day culture (**Fig. S1J**). Further, flow cytometry analysis did not reveal any significant impact of FKBP2 deletion on the expression of pluripotency markers OCT3/4 and NANOG (**Fig. S1K**). Together, we concluded the successful generation of FKBP2 KO hPSCs.

Next, we analyzed the impact of FKBP2 on insulin processing by examining the levels of proinsulin and insulin in WT and KO SC-β cells. Given the enzymatic properties of FKBP2, in KO we anticipated the increase in unprocessed proinsulin and decrease in mature insulin levels. Immunofluorescence staining showed a 70% reduction in insulin, but not proinsulin levels, in FKBP2 KO SC-β cells. Additionally, we observed the 64% elevated C-PEP expression in KO cells (**Fig. 1H-I**). Next, we utilized the Tet-On system, integrating a reverse tetracycline-controlled transactivator into the hPSC genome via piggyBac transposon, to induce ectopic expression of FKBP2-FLAG in the presence of doxycycline (**Fig. 1K** and **S1M-N**). Doxycycline treatment at the EP stage of hPSC differentiation led to an increased expression of CHGA, C-PEP, and insulin at day 8 post-treatment, representing the SC-β cell stage (**Fig. 1J-K** and **Fig. S1O**). Consistent with impaired proinsulin processing, a quantitative assessment of total insulin secretion following KCl-induced membrane depolarization demonstrated a 42% decrease in insulin production in KO cells (**Fig. 1L**). To investigate the abundance and characteristics of insulin granules, we employed structured illumination microscopy (SIM). Co-staining of insulin and vimentin (VIM) revealed a significant, yet slight (by 15%) increase in individual insulin granule size in KO compared to WT SC-β cells (**Fig. 1M**). Transmission electron microscopy (TEM) and subsequent morphometric analysis revealed that FKBP2 KO cells had an increased presence of more intensely stained insulin granules with greater size heterogeneity compared to WT SC-β cells, suggesting defects in insulin granule formation and maturation (**Fig. 1N**). Together, the absence of FKBP2 leads to a build-up of immature insulin granules, likely due to the accumulation of misfolded or unprocessed proinsulin and C-PEP aggregates, and a subsequent deficit in insulin secretion, directly linking its ER chaperone function to the integrity of the insulin secretory pathway and β cell proteostasis. The lack of concurrent proinsulin accumulation in the absence of FKBP2 might suggest protective measures such as activation of unfolded protein response (UPR) and downregulation of proinsulin synthesis, efficient ER-Associated Degradation (ERAD) actions, technical limitations of proinsulin detection methodology, or reduced SC-β cell formation.

### Single-cell RNA sequencing reveals enhanced early SC-α cell formation, reduced progenitor pool, and dysregulation of insulin processing in FKBP2-deficient cells

To verify and further investigate the changes in FKBP2 KO SC-β cells, we performed the scRNA-seq of WT and KO cells at day 31 of pancreatic differentiation. Uniform manifold approximation and projection (UMAP) plots showed a similar distribution of WT and KO cells (**Fig. S2A**). The Seurat-based analysis of sequenced 5,188 WT and 5,800 KO cells revealed the presence of 7 distinct clusters in both WT and KO samples (**Fig. 2A**). Identity of each cluster was established based on the expression of a cohort of cell type-specific marker expression, including *INS* and *NKX6-1* for SC-β cells, *GCG* and *ARX* for SC-α, *CHGA*, *HHXE* and *SST* for SC-δ, *CHGA*, *ETV1* and *NKX6-1* for endocrine progenitors 1 (EP1), *CHGA*, *NKX6-1* and *FEV* for endocrine progenitors 2 (EP2), *PDX1* and *SOX9* for pancreatic progenitors 2 (PP2), and *PDX1* and *MKI67* for pancreatic progenitors 1 (PP1) (**Fig. 2B** and **S2B**). Interestingly, we observed differential abundance of various pancreatic cell populations between WT and KO samples. Specifically, in KO samples, PP2 and EP2 populations were significantly reduced, by 0.65-fold and 0.69-fold, respectively. Conversely, EP1 and early SC-α cell populations were both significantly enriched by ∼2-fold (**Fig. 2C** and **S2B**). The KO exhibited differential expression of a cohort of genes, including upregulated α cell markers *GCG*, *ARX*, and *ETV1* as well as *ADCY9* and *FOXP2,* and downregulated expression of *HDAC9*, *INS*, and *CALY* (**Fig. 2D** and **S2C**). Pseudotime trajectory analysis using the Monocle3 algorithm ^14–17^ revealed a clear and expected developmental progression in WT cells, from PP1, PP2, having lower pseudotime values, through EP1, EP2 and then terminally differentiated cells (SC-β, SC-α, SC-δ) reaching higher pseudotime values. In contrast, FKBP2 KO cells showed a distinct developmental dynamic. While early progenitor populations (PP1, PP2, EP1, EP2) had distributions similar to WT cells, the maturation of terminal cell types was altered. The SC-β cell population in KO samples showed a slightly lower density and less spread into higher pseudotime values, indicating fewer cells reached a fully differentiated β cell state. Conversely, the SC-α population was more abundant and had a broader distribution of higher pseudotime values, suggesting that a larger proportion of KO cells were preferentially directed toward the α cell lineage (**Fig. S2D-E**). Together, suggesting distinct differentiation dynamics in KO cells compared to WT cells. Collectively, these findings demonstrate that the absence of FKBP2 disrupts the normal developmental trajectory, leading to a shift in cell fate dynamics.

**Figure 2:**
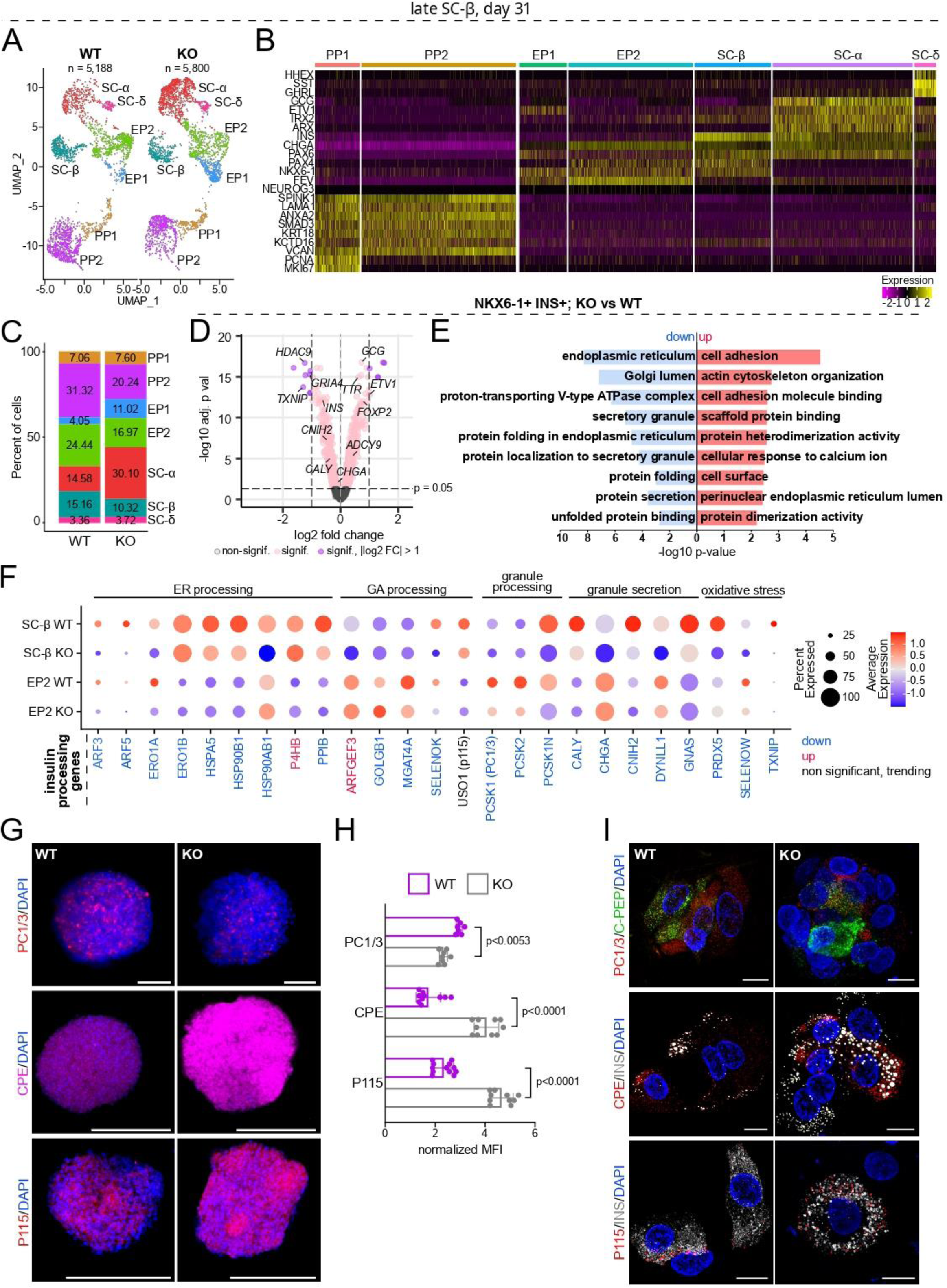
Single-cell transcriptomic analysis reveals elevated ER stress and dysregulation of insulin processing pathways in FKBP2 KO SC-β cells. **(A)** UMAP plot demonstrates main cell types at the late SC-β cell stage of differentiation (day 31), distributed into 7 different clusters, including pancreatic progenitors 1 and 2 (PP1, PP2), endocrine progenitors 1 and 2 (EP1, EP2), SC-β, SC-α, and SC-δ cells. Each dot represents a single cell, and each cell type is marked with a distinct color. **(B)** Heatmap of normalized mRNA expression of key differentiation and cell type-specific marker genes across identified clusters based on scRNA-seq of day 31 WT SC-β cells. The color intensity indicates the normalized gene expression level from −2 to 2. **(C)** Quantification of pancreatic cell populations (as % of total cells) in WT and KO samples. The bar graph illustrates the proportional representation of different cell clusters (PP1, PP2, EP1, EP2, SC-β, SC-α, SC-δ) in WT and KO conditions. **(D)** Volcano plot showing differentially expressed genes between KO and WT NKX6-1+/INS+ cells at day 31 of pancreatic differentiation. Selected genes that are significantly up- or downregulated between KO and WT endocrine cells are marked. The x-axis represents log2 fold change, and the y-axis represents -log10 adjusted p-value. Horizontal dashed line demarks p-value 0.05, while vertical dashed line marks -/+ 1.0 log2 fold change. **(E)** Gene Ontology (GO) enrichment analysis of differentially expressed genes in FKBP2 KO NKX6-1+/INS+ cells. The panel shows significantly enriched Gene Ontology terms, categorized by biological processes and molecular functions in upregulated (red) and downregulated (blue) genes from panel D. **(F)** Dot plot shows expression levels and percentage of cells expressing key genes involved in ER processing, Golgi processing, granule processing, granule secretion, oxidative stress response, and insulin processing pathways in WT and KO EP2 and SC-β cells. Dot size represents the percentage of cells expressing each gene, and color intensity represents the average expression level. **(G)** Representative immunofluorescence images of WT and FKBP2 KO SC-β cells stained for insulin processing enzymes: proinsulin convertase (PC1/3, red), carboxypeptidase E (CPE, red), and P115 (carboxypeptidase E, red). DAPI marks nuclei (blue). Scale bar, 100 µm. **(H)** The bar graph shows quantification of mean fluorescence intensity (MFI) for PC1/3, CPE, and P115 staining of WT and FKBP2 KO SC-β cells. The mean ± SD is shown, and each dot represents a single image. N = 3 independent biological repeats. An unpaired t-test was used for statistical analysis. **(I)** Immunofluorescence staining and high-resolution SIM show co-localization of C-PEP (green) with PC1/3 (red), insulin (white/gray) with CPE (red), and insulin (white/gray) with P115 (red) in WT and KO SC-β cells, at day 31. Scale bar, 10 µm.

We then focused on the *NKX6-1+/INS+* SC-β cell population, which was 1.5-fold reduced in KO samples (**Fig. 2C** and **S2B**). Gene Ontology (GO) and Kyoto Encyclopedia of Genes and Genomes (KEGG) pathway analysis of the differentially expressed genes (DEGs) in KO SC-β cells revealed the downregulation of terms associated with the ER lumen, Golgi apparatus, protein folding, protein folding in the ER, unfolded protein binding, and secretory granules (**Fig. 2E**). Consistent with FKBP2 known function as an ER chaperone involved in proinsulin folding and processing, and defective proinsulin folding. Interestingly, terms such as cell adhesion, actin cytoskeleton organization, cell surface, and cellular response to calcium ions were upregulated in KO versus WT SC-β cells (**Fig. 2E**). Analysis of the genes contributing to the changes in GO and KEGG terms revealed upregulation of *RHOB*, *ROBO1*, *NLGN1*, and *EDIL3*, genes associated with cell adhesion, as well as *NLGN1*, *ADCY8*, and *ADGRV1*, which are involved in the cellular response to calcium ion, in KO. Conversely, downregulated genes included *KCNN3*, *INS*, and *GNAS*, within the protein secretion term, and *PPIB*, *ERP29*, *HSP90B1*, *HSPA5*, and *PDIA3*, which are key mediators of protein folding (**Supplementary Table 5**). Moreover, *INS+/NKX6-1+* SC-β cells exhibited lower expression levels of ER processing-related genes like *ERO1B*, *HSPA5,* and *PPIB*, as well as Golgi apparatus-associated genes, including *GOLGIB1* and *MGAT4A,* following FKBP2 deletion (**Fig. 2F** and **S2F**). Finally, genes linked to granule processing (*PCSK1*, *PCSK2*), granule secretion (*CHGA*, *AMPH*, *DYNLL1,* and *GNAS*), and oxidative stress (*TXNIP*, *PRDX5,* and *SELENOW*) were downregulated in KO EP2 and SC-β cells (**Fig. 2F** and **S2F**). We then confirmed changes in selected gene expression at the protein level in KO SC-β cells. Immunofluorescence combined with confocal microscopy and quantification revealed that two key enzymes involved in proinsulin post-translational processing were dysregulated in KO SC-β cells. Proinsulin convertase 1/3 (PC1/3), responsible for initiating proinsulin cleavage, showed 1.35-fold decreased protein levels (**Fig. 2G-I**). Conversely, carboxypeptidase E (CPE), which completes insulin maturation by removing basic amino acids after PC1/3 cleavage, exhibited 2.3-fold increased protein expression (**Fig. 2G-I** and **S2G**). This increase in CPE is likely a compensatory response to inefficient proinsulin processing in KO cells. Finally, p115, a vesicular transport factor that plays a crucial role in the biogenesis and maintenance of the Golgi apparatus, as well as in transport between the ER and Golgi ^18^ was increased in KO SC-β cells (**Fig. 2G-I**). Together, these findings suggest defective proinsulin processing and insulin granule maturation in the absence of FKBP2 in SC-β cells.

### Elevated ER stress in FKBP2-deficient SC-β cells

Next, we investigated whether loss of ER chaperone and deficient proinsulin processing lead to increased ER stress in KO SC-β cells. Immunofluorescence and quantification of key indicators of the UPR activation ^19^, showed increased protein expression of PKR-like ER kinase (PERK) by 2-fold and Activating Transcription Factor 6 (ATF6) by 1.4-fold (**Fig. 3A**). While the expression of genes associated with oxidative stress, including *PRDX5*, *SELENOW,* and *TXNIP,* was decreased in KO SC-β cells (**Fig. 2F** and **S2F**), which might be an initial, adaptive response of cells to the misfolded proinsulin accumulation and severe ER stress ^20,21^. The ER functions as the primary intracellular calcium storage, and elevated ER stress can lead to an increase in cytosolic Ca^2+^ concentration through several possible mechanisms, including compromised Sarco/Endoplasmic Reticulum Calcium ATPase (SERCA) function and ER membrane “leakiness” ^22^. Immunofluorescence combined with confocal microscopy and quantification revealed a decreased protein expression of SERCA2, an ATP-dependent Ca²⁺ pump and ER marker ^23,24^ (**Fig. 3B** and **C**). Consistent with elevated ER stress and reduced SERCA2 expression, we observed increased intracellular Ca^2+^ levels in KO SC-β cells (**Fig. 3D**).

**Figure 3:**
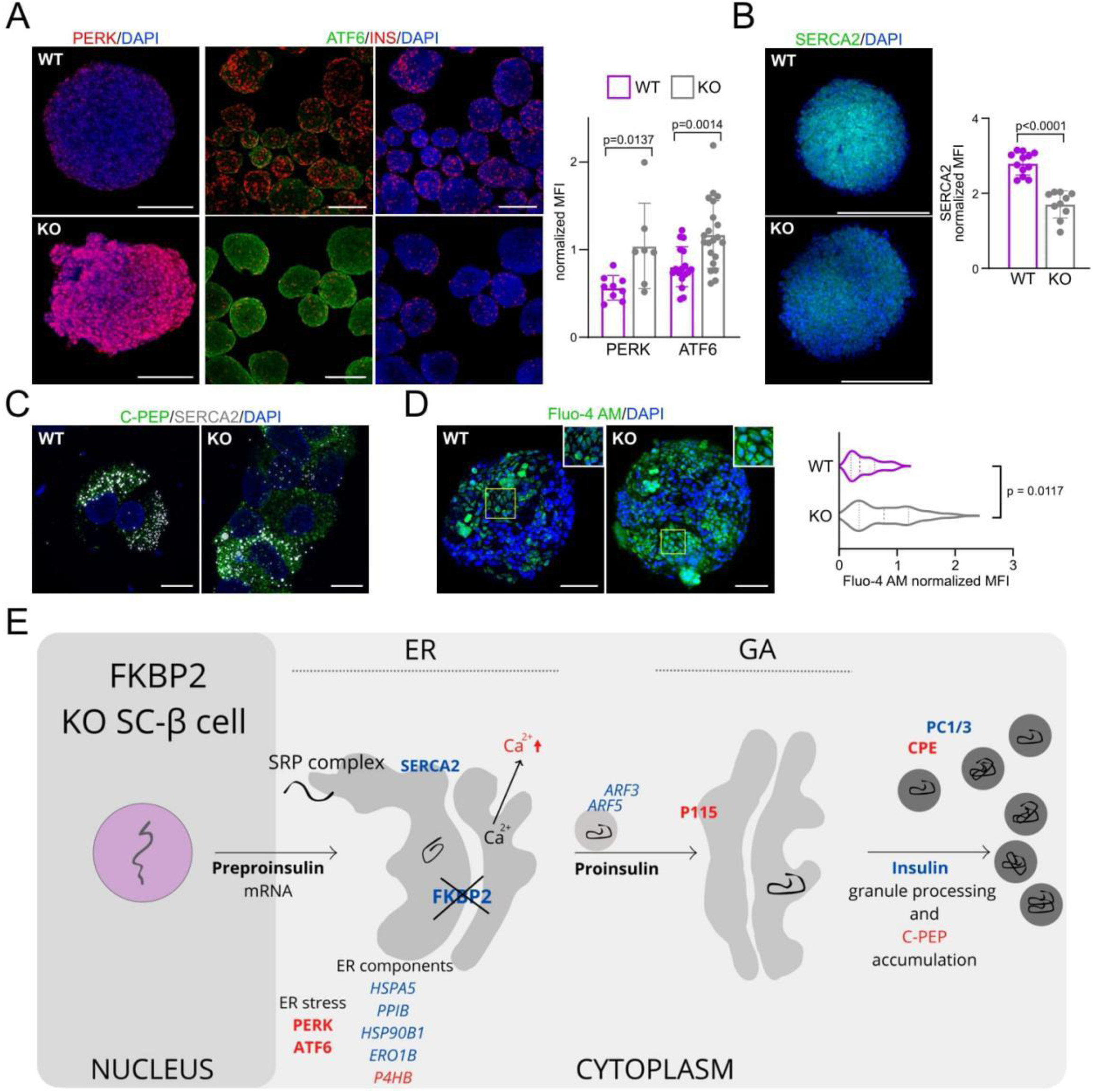
FKBP2 loss triggers ER stress response, leading to intracellular calcium increase. **(A)** Immunofluorescence staining shows expression of PERK (red) and ATF6 (green) with insulin (INS, red) in WT and FKBP2 KO SC-β cells. DAPI marks nuclei (blue). The bar graph below shows the quantification of the mean fluorescent intensity (MFI) for PERK and ATF6 staining in WT and KO conditions. The mean ± SD is shown, and each dot represents a single image. N = 3 independent experimental repeats. An unpaired t-test was used for statistical analysis. Scale bar, 100 µm. **(B)** Immunofluorescence staining of WT and FKBP2 KO SC-β cells shows SERCA2 (green) expression. DAPI marks nuclei (blue). Scale bar, 100 µm. On the right, the bar graph shows the quantification of the mean fluorescent intensity (MFI) for SERCA2 in WT and KO conditions. Mean ± SD is shown, and each dot represents a single image. N = 3 independent biological repeats. An unpaired t-test was used for statistical analysis. **(C)** Immunofluorescence staining and high-resolution SIM show co-localization of C-PEP (green) with SERCA2 (white/gray) in WT and KO SC-β cells. Scale bar, 10 µm. **(D)** Immunofluorescence and quantification of cytoplasmic calcium levels. Cells were stained with the calcium indicator Fluo-4 AM (green) and DAPI (blue). Scale bar, 100 µm. Insets show magnified views of the highlighted areas. The violin plot below displays the distribution of Fluo-4 AM mean fluorescence intensity (MFI) normalized to DAPI for WT and KO cells. P-values were calculated using a paired Student t-test. The violin plots show the data distribution with the median indicated as a dashed line and IQR as a dotted line, representing 25th–75th percentiles. N = 3 independent experiments is shown. **(E)** Proposed model of the role of FKBP2 in proinsulin processing and ER homeostasis in SC-β cells. In the absence of FKBP2, ER components, including SERCA2, are reduced. FKBP2 acts as a chaperone that facilitates the proper folding of proinsulin within the ER. Without FKBP2, the ER undergoes stress, leading to the upregulation of ER stress markers (PERK and ATF6). The disrupted ER environment and reduced SERCA2 levels lead to elevated cytoplasmic Ca^2+^ and impaired proinsulin folding. This results in the accumulation of misfolded proinsulin, leading to a downstream failure in the processing of proinsulin into mature insulin and C-PEP, which ultimately impacts insulin granule formation and secretion. Altered expression is observed in key genes involved in these processes, including ER-related (HSPA5, PPIB, HSP90B1, ERO1B, P4HB), involved in trafficking (ARF3, ARF5, USO1/P115), and PROINS processing protein (PCSK1/PC1/3, CPE). Protein symbols are written in uppercase regular letters, whereas gene symbols are written in uppercase italics. Genes and proteins marked in blue are downregulated, whereas marked in red are upregulated.

Collectively, these data suggest that FKBP2 absence initiates a critical cascade of events. The misfolding of proinsulin, triggered by the lack of FKBP2 isomerase activity, is the primary event. The misfolded proinsulin then activates the UPR, leading to a self-reinforcing cycle where intensified ER stress and elevated intracellular calcium promote further protein misfolding. Concurrently, the deletion of FKBP2, given its additional role as an ER chaperone, also contributes to disturbed ER homeostasis. These widespread cellular disruptions, encompassing compromised ER homeostasis, impaired ER-Golgi apparatus transport, and altered calcium signaling, collectively contribute to defective granule insulin formation, culminating in severe ER stress and significantly reduced insulin secretion (**Fig. 3E**).

### Altered pancreatic endocrine cell development following FKBP2 deletion

Our scRNA data suggested that the absence of FKBP2 might alter endocrine differentiation. We, therefore, investigated whether FKBP2 regulates SC-β formation. In humans, EPs and endocrine cells start to form between 8 and 13 post-conception weeks (PCW) of development. Based on scRNA-seq data of the human pancreas during embryogenesis, we observed an increase in *FKBP2* mRNA expression from PCW 4 to 11 in progenitors and endocrine cells (**Fig. 4A** and **S3A-C**). Immunofluorescence staining of human pancreas at PCW13 revealed FKBP2 expression in the developing pancreatic epithelium. Cells positive for C-PEP were detected in close proximity to the pancreatic epithelium, consistent with the detachment of β cells from the epithelium during development. Most C-PEP+ cells co-expressed FKBP2, although not all FKBP2+ cells were C-PEP+, suggesting FKBP2 present in CHGA+ endocrine progenitors or other endocrine cell types (**Fig. 4B**).

**Figure 4:**
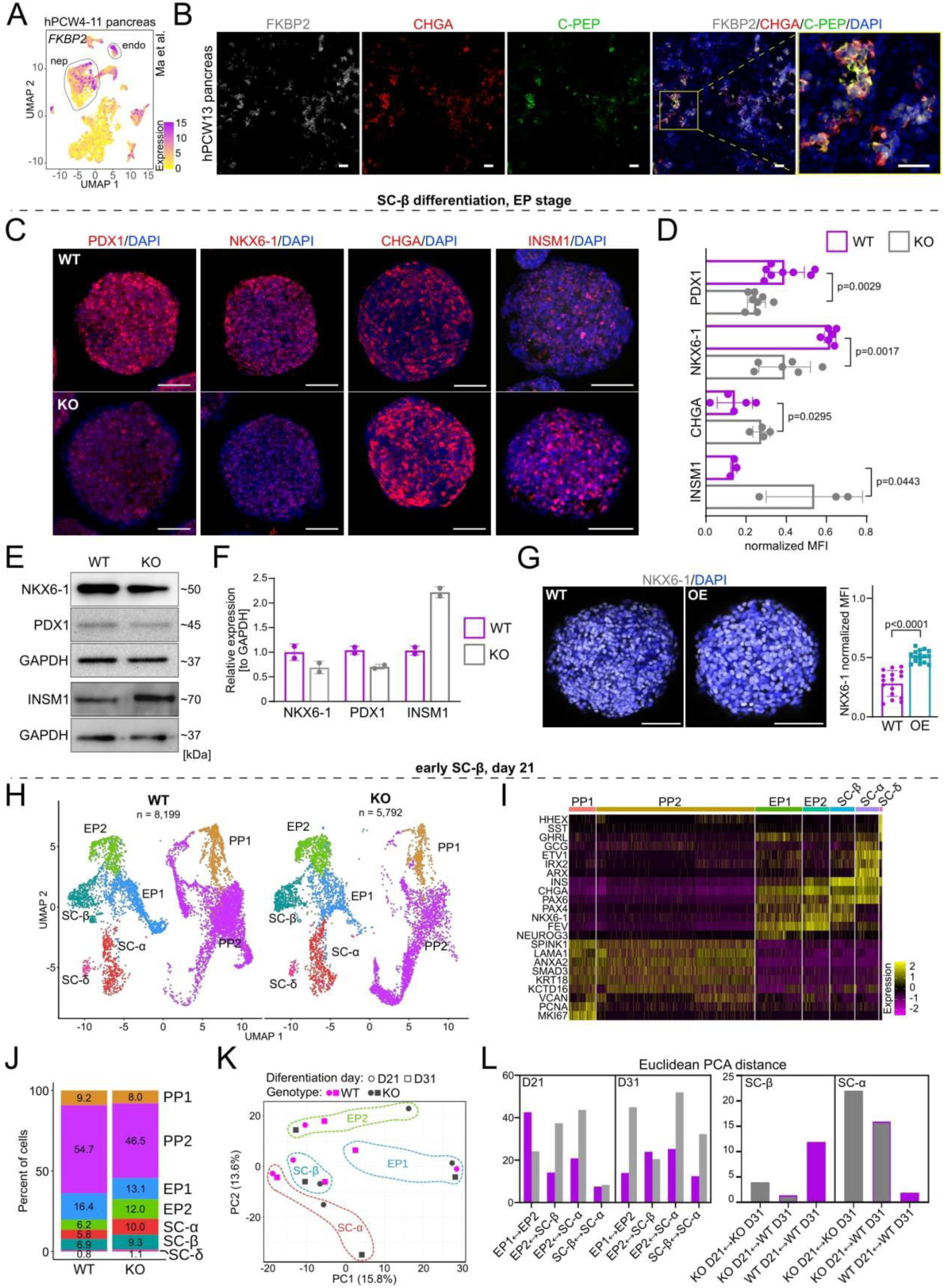
Loss of FKBP2 impairs differentiation of pancreatic endocrine cells. **(A)** Uniform Manifold Approximation and Projection (UMAP) plot showing pancreatic cells from the hPCW4-11 human fetal pancreas ^42^ with *FKBP2* expression. Endocrine (endo) and non-endocrine/pancreas (nep) cell populations are indicated by labeled regions and dashed outlines. **(B)** Immunofluorescence images of human fetal pancreas at post-conception week 13 (PCW13) stained for FKBP2 (gray/white), CHGA (red), and C-peptide (green) expression. DAPI marks nuclei (blue). The rightmost panel displays a merged image of FKBP2, CHGA, C-PEP, and DAPI highlighting the co-localization of FKBP2 with endocrine cell markers. Scale bar, 100 µm. **(C)** Immunofluorescence staining of PDX1, NKX6-1, CHGA, and INSM1 (all red) in WT and FKBP2 KO SC-β cells. DAPI marks nuclei (blue). Scale bar, 100 µm. **(D)** Bar plot shows the quantification of mean fluorescent intensity (MFI) for PDX1, NKX6-1, CHGA, and INSM1 in WT and KO cells at the EP stage. Mean ± SD is shown, with each dot on the graph representing a single image, N = 3-4 independent biological repeats. An unpaired t-test was used for statistical analysis. **(E)** Western blot analysis of NKX6-1, PDX1, and INSM1 in WT and FKBP2 KO EPs. GAPDH used as a loading control. Representative blots from 2 biological replicates are shown. **(F)** Relative expression levels of NKX6-1, PDX1, and INSM1 normalized to GAPDH. N = 2 biological repeats. **(G)** Representative immunofluorescent staining shows NKX6-1 expression in WT and FKBP2-OE cells at day 26 of differentiation. OE was induced on day 20. Scale bar, 100 µm. The bar graph, on the right, shows quantification of mean fluorescent intensity (MFI) for NKX6-1 in WT and OE cells. Mean ± SD is shown, and each dot on the graph represents a single image. N = 3 independent biological repeats, and an unpaired t-test was used for statistical analysis. **(H)** UMAP visualization based on scRNA-seq of WT and FKBP2 KO cells at day 21 reveals different cell clusters, including pancreatic progenitors (PP1, PP2), endocrine progenitors (EP1, EP2), and more mature cell types (SC-α, SC-β, SC-δ). **(I)** Heatmap of normalized mRNA expression of key marker genes across distinct clusters based on scRNA-seq of day 21 WT SC-β cells. The color intensity indicates the expression level from - 2 to 2. **(J)** Stacked bar chart showing the percentage of cells in each cluster (PP1, PP2, EP1, EP2, SC-α, SC-β, and SC-δ) for WT and KO samples at day 21 of pancreatic *in vitro* differentiation. **(K)** Principal component analysis (PCA) plot shows the distribution of cell clusters (EP1, EP2, SC-β, SC-α) at different differentiation days (D21 and D31) for WT and KO genotypes. **(L)** Bar graphs comparing Euclidean PCA distances between cell clusters at D21 and D31 for WT and KO cells. The distances reveal differences in the differentiation trajectories between WT and KO cells. For example, the distance between the EP2 and SC-α clusters is larger in KO cells, indicating altered progression towards the endocrine cell fate.

We next examined the impact of FKBP2 loss on *in vitro* β cell differentiation. At the PP stage, we did not observe significant differences in the expression of NKX6-1, a transcription factor crucial for pancreatic progenitor development (**Fig. S3D**). However, 8 days later, at the EP stage, FKBP2 KO resulted in a significant decrease in protein levels of PDX1 and NKX6-1, and increased expression of endocrine cell markers, including CHGA and insulinoma (INSM1) (**Fig. 4C** and **D**). The western blot analysis further corroborated the lower PDX1 and NKX6-1, along with higher INSM1 expression in KO EPs (**Fig. 4E** and **F**). Moreover, upon FKBP2 OE at EP cell stage, we observed the opposite effect, the increase in NKX6-1 protein level (**Fig. 4G**).

Given the FKBP2 presence in EPs and newly formed endocrine cells, changes in KO transcriptomes revealed by scRNA-seq at day 31, and with immunofluorescence-based manifestation of altered differentiation of FKBP2 KO cells, we next aimed to further investigate the differentiation defect in FKBP2 KO endocrine cells. To this end, we performed the scRNA-seq of 8,199 WT and 5,792 KO cells at day 21 of pancreatic differentiation (**Fig. 4H** and **S4A**). Seurat analysis separated WT and KO cells into 7 clusters, which, based on the expression of well-established markers, were assigned as PP1, PP2, EP1, EP2, SC-β cells, SC-α cells, and SC-δ cells **(Fig. 4I, S4C,** and **Supplementary Table 1**). Interestingly, we observed a 1.93-fold increase in the presence of more differentiated progenitors, EP2, and all endocrine cell types in KO samples compared to the WT population, suggesting accelerated differentiation in the absence of FKBP2 (**Fig. 4J** and **S4B**). Among endocrine cells, the highest increase by 1.72-fold was observed for SC-α cells (**Fig. 4J** and **S4B**). Corresponding with changes in cluster frequency, gene expression analysis revealed altered expression of multiple endocrine markers, including *NKX6-, FEV, CHGA* and *GCG* (**Fig. S4C**).

To investigate how FKBP2 deletion alters differentiation kinetics, we merged the scRNA-seq datasets for day 21 and day 31 WT and KO cells and performed the principal component analysis (PCA) analysis. Principal Component 1 (PC1) accounts for 15.8% of the variance and largely separates the cells by their differentiation day, while Principal Component 2 (PC2) accounts for 13.6% of the variance and mainly distinguishes between the different cell populations. The PCA plot illustrated a clear trajectory of differentiation from earlier stages (EP1, EP2) to later stages (SC-β, SC-α) and showed that the KO cells exhibit a different distribution at both day 21 and day 31 compared to the WT cells, suggesting an altered differentiation pathway or kinetics. Specifically, KO cells at day 21 are more spread out on the PCA plot compared to WT cells, indicating a less uniform population (**Fig. 4K**). This observation aligns with the shifts in population percentages (**Fig. 4J**), which suggests that the absence of FKBP2 disrupts the normal progression toward SC-β cells.

Interestingly, the day 31 KO SC-β cells clustered closely with the day 21 WT SC-β cells, indicating a more advanced differentiation state upon FKBP2 loss (**Fig. 4K**). Further analysis of both Euclidean ^25^ and Mahalanobis ^26^ distances supported a more distinct and variable differentiation process in KO cells. At day 21, Euclidean distances are lower (e.g., ∼20 for EP1+EP2+SC-α+SC-β), increasing at day 31 (e.g., ∼40–50), indicating growing separation as differentiation progresses. Higher values suggest greater differences in the PCA space, reflecting changes in cell population structure or gene expression profiles. We have also calculated the Mahalanobis distances, which adjust for covariance, providing a statistically robust measure of separation that accounts for variable correlations. In general, Mahalanobis distances mirror the Euclidean increase, with day 31 showing higher separation (e.g., ∼4 for EP1+EP2+SC-α+SC-β vs. ∼1-3 for day 21). The Mahalanobis distance for KO SC-β cells at day 21 was smaller when compared to WT cells at day 31 than when comparing WT cells at day 21 and 31. This suggests that KO SC-β cells at day 21 were more similar to WT SC-β cells at day 31 than to their WT counterparts at day 21, indicating a more rapid differentiation trajectory. Conversely, the opposite trend was observed for SC-α cells (**Fig. 4L** and **S4D**).

Collectively, these scRNA-seq analyses demonstrate that FKBP2 deletion disrupts the differentiation program, altering not only the location of cells in PCA space but also their variance and covariance patterns within respective clusters.

### FKBP2 regulates α cell vs β cell fate *in vitro* determination

The scRNA at day 21 and 31 revealed distinct alterations in endocrine cell populations between WT and KO cells. At day 21, the ratio of KO to WT cells in the SC-β cluster was approximately 1.4, indicating a slight increase in β cell formation in the absence of FKBP2. However, this trend was reversed at day 31, and the KO/WT ratio for SC-β was 0.8. Conversely, the SC-α cell cluster showed an increased KO/WT ratio of approximately 1.8 at day 21 and 2.2 at day 31, suggesting enhanced SC-α cell differentiation at both time points (**Fig. 5A**). Expression analysis of key pancreatic hormones confirmed significant dysregulation in FKBP2 KO differentiation. Insulin expression was initially increased (at day 21) but then reduced in KO cells compared to WT at day 31, consistent with altered β cell differentiation. In contrast, GCG expression was markedly elevated in KO cells at both time points, supporting the observed shift toward α cell fate (**Fig. 5B**).

**Figure 5.**
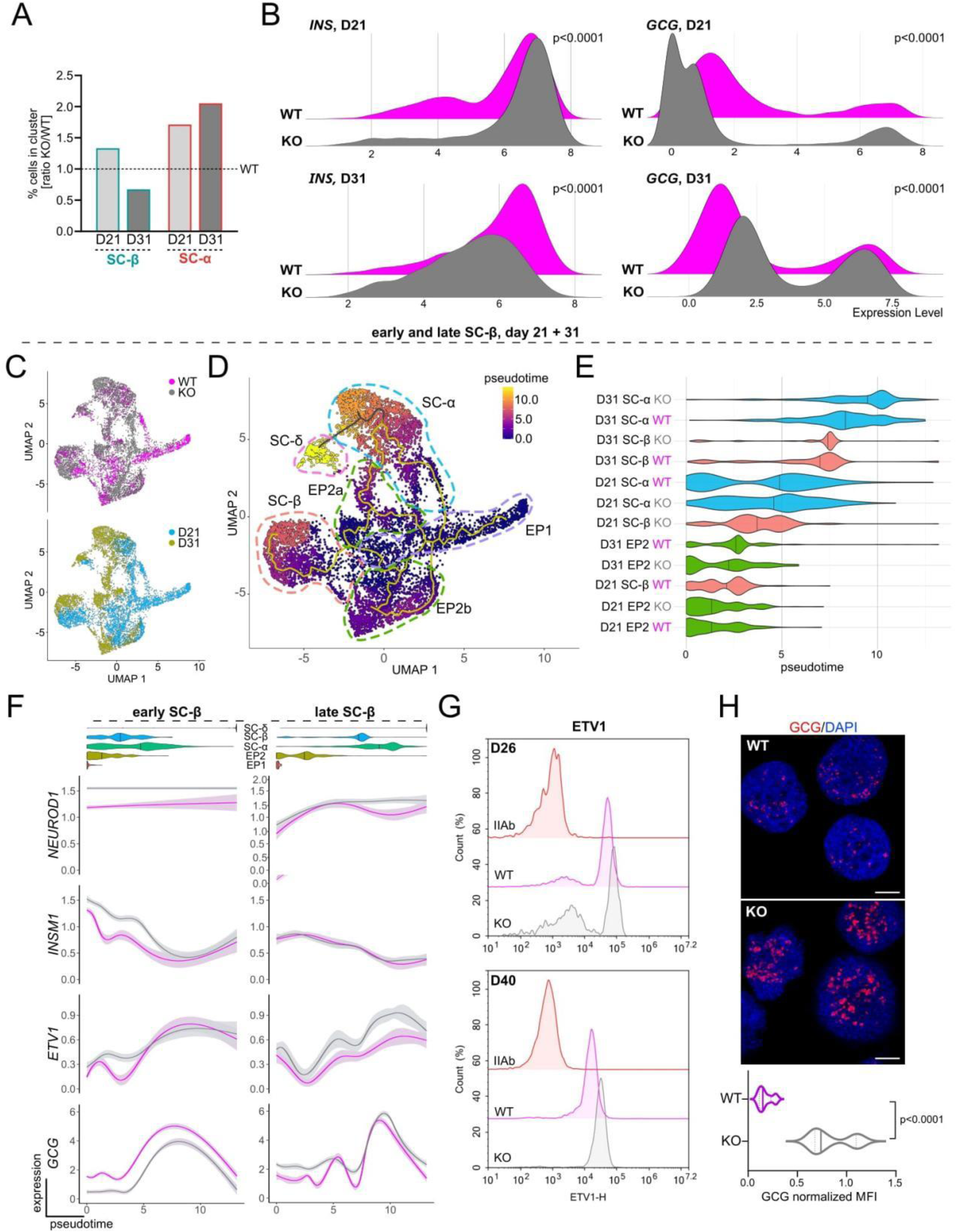
Developmental trajectory analysis and glucagon expression in WT and FKBP2 KO pancreatic endocrine differentiation. **(A)** Bar plot shows the ratio of KO to WT cell percentages of INS+/NKX6-1+ SC-β cells and GCG+ SC-α at day 21 (D21) and day 31 (D31) of differentiation in WT and FKBP2 KO hPSCs. **(B)** Ridge plots comparing expression of insulin (INS) and glucagon (GCG) at day 21 and day 31 of differentiation in WT and FKBP2 KO hPSCs. **(C)** UMAP visualization showing the distribution of WT and KO cells (top) and visualization of early (day 21) and late (day 31) SC-β cells (bottom) during pancreatic differentiation. **(D)** UMAP visualization of early (day 21) and late (day 31) SC-β cells showing clustering of endocrine cell populations and pseudotime trajectory analysis. Cells are colored by pseudotime values (0-10 scale) and annotated with cell type identities: endocrine progenitor 2a and 2b (EP2a, EP2b), stem cell-derived α cells (SC-α), β cells (SC-β), and δ cells (SC-δ). **(E)** Violin plots showing pseudotime distributions of WT and KO cells across different endocrine populations at D21 and D31. **(F)** Expression dynamics along pseudotime trajectories for key transcription factors and hormones in both early and late SC-β cell differentiation. Plots show expression levels of *GCG*, *ETV1*, *INSM1,* and *NEUROD1* across pseudotime for WT and KO conditions, with separate trajectories indicated for early and late SC-β cell populations. **(G)** Flow cytometry analysis of ETV1 protein expression in WT and KO cells at day 26 (D26) and day 40 (D40) of differentiation. IIAb staining serves as control. **(H)** Representative immunofluorescence images of glucagon (GCG, red) with DAPI nuclear staining (blue) in WT and KO conditions, showing cellular localization and expression patterns. Scale bar, 100 µm. Quantification of normalized mean fluorescence intensity (MFI) for GCG expression comparing WT and KO conditions. The violin plot below shows the data distribution with the median shown as a dashed line and IQR as a dotted line, 25th–75th percentiles. N = 3 independent biological repeats. An unpaired t-test was used for statistical analysis.

We then used pseudotime trajectory analysis to gain insights into the differentiation dynamics of endocrine cell specification in absence of FKBP2. UMAP visualization of early and late SC-β cells (day 21, day31) revealed proper developmental trajectories spanning EP1, EP2a, EP2b, and terminally differentiated SC-α, SC-β, and SC-δ cells in both conditions (**Fig. 5C** and **D**). Pseudotime analysis demonstrated no major differences in EP2 populations at either day 21 or day 31. At day 21, KO cells showed a relative enrichment of SC-β compared to WT, consistent with accelerated β cell differentiation. By contrast, at day 31 this trend was reversed, with KO displaying a predominance of SC-α cells, indicating a shift in lineage allocation from β to α cell fate. Other populations showed no significant differences (**Fig. 5E**). Expression dynamics along the pseudotime trajectories revealed differential regulation of key transcription factors and hormones. The expression of *NGN3* targets, *NEUROD1* and *INSM1,* was initially (at day 21) increased in the KO cells, but by day 31, it was similar to that of WT cells. The mRNA level of *ETV1*, a gene associated with EP differentiation ^27^ and α cell marker ^9,28^, was increased at day 21 and 31 in KO cells. Consistently, *GCG* expression showed a sustained increase in KO throughout both early and late developmental trajectories (**Fig. 5F**). These findings suggest that FKBP2 KO disrupts normal transcriptional programs governing endocrine cell fate specification. Immunofluorescence analysis confirmed the select scRNA-seq findings at the protein level. Flow cytometry analysis of WT and KO cells at day 40 of differentiation showed higher ETV1 protein levels in KO cells (**Fig. 5G**), while immunostaining revealed significantly increased GCG expression following FKBP2 deletion compared to WT (**Fig. 5H**).

Pseudotime analysis revealed that the *FKBP2* deletion disrupts the transcriptional dynamics of SC-β cells in a stage-dependent manner, leading to an impaired differentiation trajectory (**Fig. S5A**). During the early SC-β stage, KO cells showed a reduced *NEUROG3* expression and a premature increase in *MAFA* and *CALY* mRNA, suggesting a dysregulation in their initial programming. At the same stage, *FEV* and *CHGA* mRNA levels were transiently elevated in KO SC-β cells in day 21 KO cells compared to WT, while *PDX1* expression was decreased and *NKX6-1* remained largely unchanged. By the late SC-β stage (day 31), these early elevations in *FEV* and *CHGA* became comparable between KO and WT, indicating a potential developmental delay or transient compensatory effect. However, KO cells progressively diverged from WT in endocrine lineage specification: *INS* expression, initially similar to WT at day 21, declined markedly in KO cells by day 31. In parallel, *MAFB* levels were reduced in SC-β KO, and *ARX* expression became specifically elevated in KO cells, indicating a bias toward an α cell fate (**Fig. S5A**). Together, these findings indicate that *FKBP2* deletion alters endocrine *in vitro* differentiation in a temporally regulated manner, blocking proper β cell programming at day 21 and enhancing α cell formation by day 31.

### HDAC9 expression is dysregulated in FKB2 KO EPs and SC-β cells

Interestingly, one of the most deregulated genes identified in scRNA-seq of KO SC-β cells is Histone Deacetylase 9 (*HDAC9*) with a 2-fold change (**Supplementary Table 2**). HDAC9, a member of the class IIa histone deacetylases, is expressed in pancreatic β cells and Hdac9^−/−^ mice exhibit increased β cell mass ^29^. At day 21 of differentiation, *HDAC9* mRNA was elevated in the EP1 and EP2 KO cells (**Fig. 6A, B**, and **Supplementary Table 3**). While at day 31, *HDAC9* mRNA was significantly downregulated in EP1 and EP2 KO, which may be part of a compensatory response following FKBP2 deletion (**Fig. 6D, E**, and **Supplementary Table 4**). In line with these findings, UMAP visualization revealed increased *HDAC9* expression in KO cells at day 21, whereas pseudotime analysis demonstrated its marked downregulation by day 31 (**Fig. S6A** and **S6B**). Immunofluorescence staining showed 3- to 2.5-fold higher HDAC9 protein levels at early (day 21) and late (day 31) β cell stage of differentiation, respectively **(Fig. 6C** and **F**). High-resolution SIM revealed nuclear localization of HDAC9 in WT and KO samples, suggesting that the increase in its expression is functionally relevant (**Fig. 6G**). To establish the link between ER stress and HDAC9 expression, we treated WT EPs with 1 µg/ml dithiothreitol (DTT), causing elevated ER stress as confirmed by increased ATF6 expression, and observed the elevated HDAC9 expression (**Fig. 6H** and **I**). Finally, to investigate the role of this increased HDAC9, we treated the KO cells at the EP stage with a well-established HDAC class IIa inhibitor, TMP195 ^30^. The six-day treatment resulted in a 2.5-fold downregulation of GCG+ cell formation and a concomitant upregulation in insulin expression (**Fig. 6J** and **K**). This is consistent with previous findings where pharmacological inhibition of class IIa HDACs, including HDAC9, by small molecule inhibitor MC1568, promoted murine *ex vivo* β cell development at the expense of other endocrine lineages ^29^. These results from independent studies reinforce the idea that class IIa HDAC activity acts as a barrier for efficient β cell differentiation, while at the same time creating a permissive environment for α cell specification. Together, we interpret these data that FKBP2 deletion at EP and early SC-β cell stages, disturbs ER homeostasis and leads to elevated ER stress, which in turn increases HDAC9 expression and causes the enhanced α cell lineage commitment.

**Figure 6:**
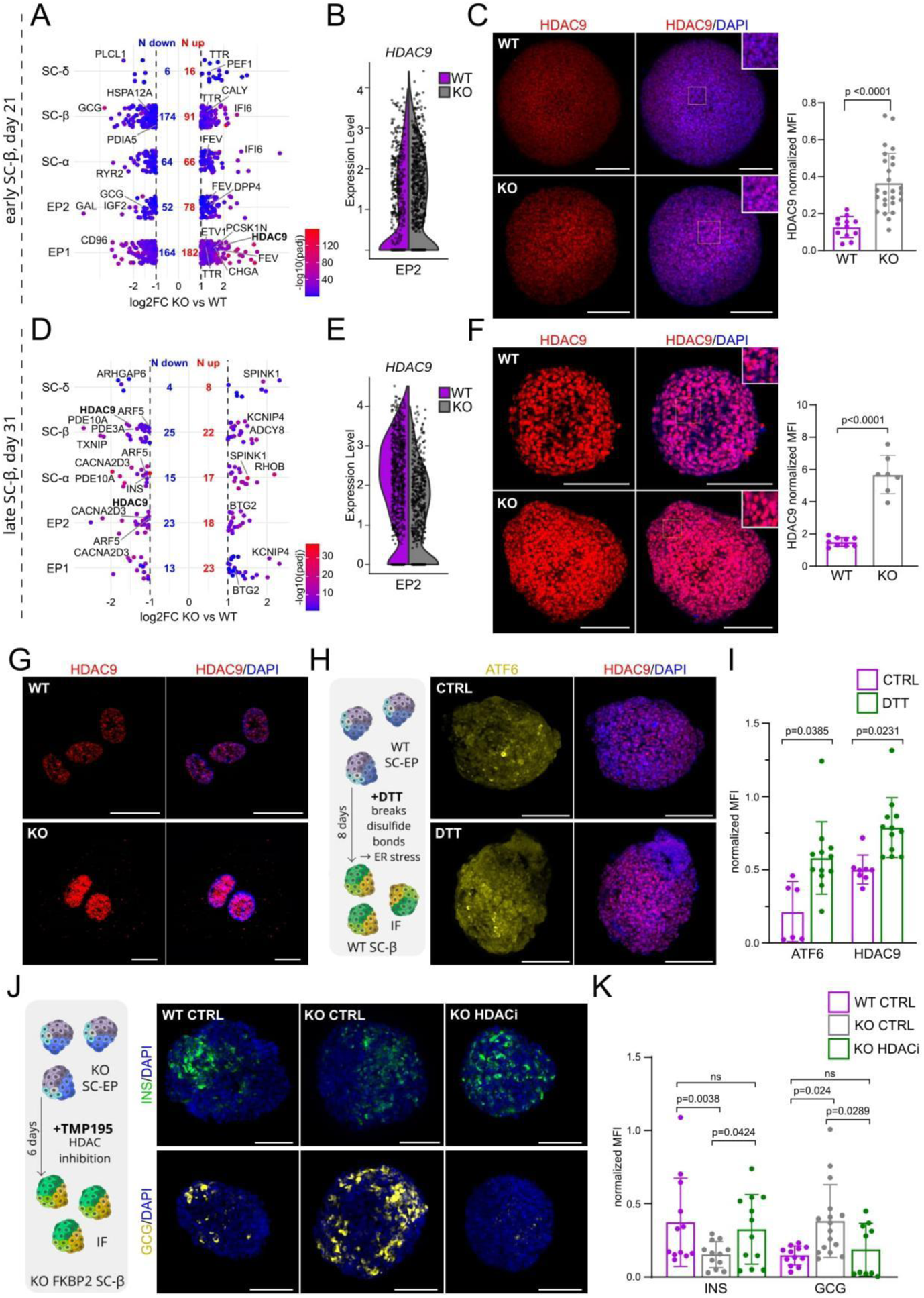
FKBP2 deletion and ER stress lead to enhanced HDAC9 expression and preferential SC-α cell formation. **(A)** Volcano plot shows differential gene expression based on scRNA-seq of early SC-β at day 21 comparing KO vs. WT conditions. Points represent individual genes with log2 fold change (x-axis) and statistical significance as -log10 (adjusted p-value) (y-axis). HDAC9 and selected differentially expressed genes are labeled. Numbers indicate genes significantly upregulated (N up) or downregulated (N down) in each cell type: endocrine progenitor 1 (EP1), endocrine progenitor 2 (EP2), stem cell-derived α cells (SC-α), β cells (SC-β), and δ cells (SC-δ). **(B)** Violin plots comparing the expression of *HDAC9* mRNA in WT and KO EP2 at day 21 of differentiation. **(C)** Representative immunofluorescence staining of β cell spheres at day 21 shows HDAC9 (red) expression in WT and KO cells. DAPI marks nuclei in blue. Scale bar, 100 µm. The graph on the right shows quantitative analysis of the normalized mean fluorescence intensity (MFI) for HDAC9 in WT and KO early SC-β cells. N = 3 biological repeats. Data are presented as mean ± SD, with each dot representing a single image. An unpaired t-test was used for statistical analysis. **(D)** Volcano plot showing differential gene expression in late SC-β cells at day 31 comparing KO versus WT conditions, with format identical to panel A. Selected genes, including HDAC9, GCG, and others, are labeled. **(E)** Violin plots comparing the expression of *HDAC9* mRNA in WT and KO EP2 at day 31 of differentiation. **(F)** Representative immunofluorescence staining of late (day 31) WT and KO SC-β cells shows HDAC9 (red) nuclear and cytoplasm localization. DAPI marks nuclei in blue. Scale bar, 100 µm. The graph on the right shows quantification of normalized HDAC9 mean fluorescent intensity (MFI) in WT and KO late (day 31) SC-β cells. N = 5 independent biological repeats. Data are presented as mean ± SD, with each dot representing a single image. An unpaired t-test was used for statistical analysis. **(G)** Representative high-resolution SIM images of SC-β cells show the localization of HDAC9 (red) and DAPI (blue) in FKBP2 KO and WT cells. Scale bar, 10 µm. **(H)** Left: experimental design to assess the impact of ER stress on HDAC9 expression in SC-β cells. WT cells at the EP stage (day 18) were treated with 1 µg/ml dithiothreitol (DTT) for 8 days to induce ER stress, and ATF6 and HDAC9 protein expression was analyzed at day 26 (SC-β cells). Right: representative immunofluorescence images of ATF6 (yellow) and HDAC9 (red) co-localization in control (CTRL) and DTT-treated WT SC-β cells. DAPI marks nuclei in blue. Scale bar, 100 µm. **(I)** Quantification of normalized ATF6 and HDAC9 mean fluorescent intensity (MFI) under CTRL and DTT treatment conditions. N = 3 independent biological repeats. Each dot represents one image; lines indicate mean ± SD. An unpaired t-test was used to assess significance, and p-values are indicated. **(J)** Left: experimental design to assess the impact of pharmacological HDAC class IIa inhibition on INS+ and GCG+ cell formation from KO hPSCs. FKBP2 KO cells at the EP stage (day 20) were treated with 1 µM TMP195 for 6 days and assessed for INS and GCG protein expression. Right: representative immunofluorescence images of INS (green) and GCG (yellow) expression in WT untreated (WT CTRL), KO untreated (KO CTRL), and KO TMP195-treated (KO HDACi) SC-β cells. DAPI marks nuclei in blue. Scale bar, 100 µm. **(K)** Quantification of normalized INS and GCG mean fluorescent intensity (MFI) in WT and KO cells, both untreated (CTRL) and treated KO with TMP195 (KO HDACi). N = 3 independent biological replicates. Each dot represents one image; lines indicate mean ± SD. An unpaired t-test was used to assess significance, and p-values are indicated.

### Increased NFAT expression in FKBP2 KO SC-β cells, possibly regulates HDAC9 expression in a calcium-dependent manner

To investigate how the loss of the ER chaperone leads to increased HDAC9 expression in KO cells, we analyzed putative transcriptional *HDAC9* regulators. Using the EnrichR TFs_TRANSFAC_JASPAR database, we identified NFATC1 (also known as NFAT2) as one of the predicted regulators and found that the *HDAC9* locus contains a putative NFAT2 binding motif. Further, motif enrichment analysis revealed that NFAT2 binding motifs were significantly less represented in KO cells (**Fig. 7A** and **B**). NFATs were shown to regulate the expression of genes involved in β cell function, including those responsible for glucose sensing and insulin secretion in mice ^31,32^.

**Figure 7:**
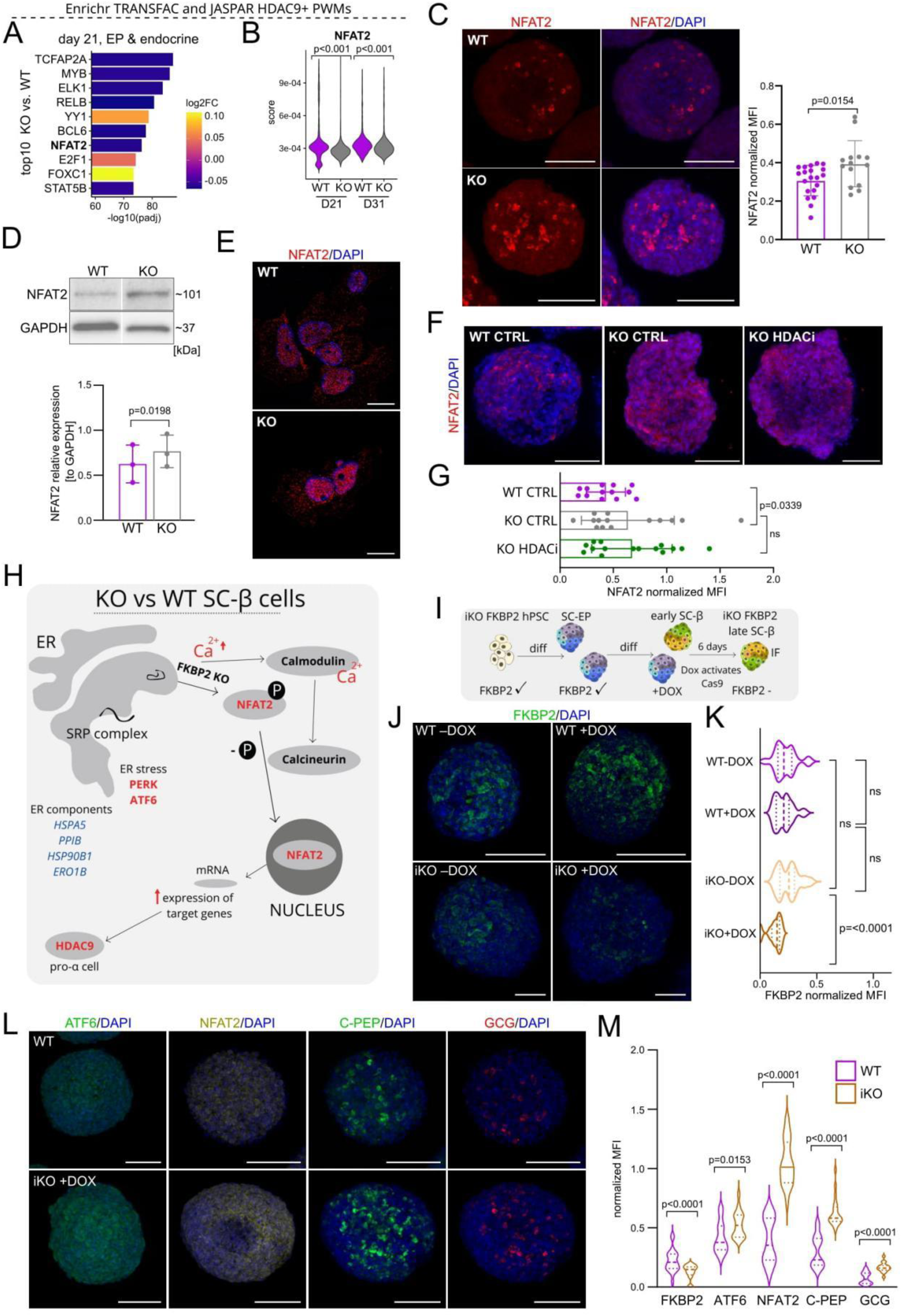
Increased NFAT expression in FKBP2-deficient endocrine cells. **(A)** Differential enrichment analysis of transcription factor (TF) binding gene sets (Enrichr TRANSFAC and JASPAR position weight matrices (PWMs) gene set filtered for TFs that have their PWMs in the HDAC9 promoter). The 10 most significant TF gene sets between KO versus WT cells at day 21 are shown. Bar colors indicate log2 fold change of gene set score between KO and WT, and the X-axis represents the -log10(padj) value. **(B)** Violin plots showing the NFAT2 gene set (Enrichr TRANSFAC and JASPAR PWMs) scores in WT and KO cells at day 21 (D21) and day 31 (D31). Statistical significance of scores difference between KO and WT cells for each day is shown by p-values (Wilcoxon rank sum test). **(C)** Immunofluorescence images show NFAT2 (red) expression in WT and FKBP2 KO cells. DAPI marks nuclei in blue. Scale bar, 100 µm. The graph on the right shows quantitative analysis of the normalized mean fluorescence intensity (MFI) for NFAT2 in WT and KO early SC-β cells. Mean ± SD from N = 3 independent biological repeats is shown, with each dot representing a single image. An unpaired t-test was used to assess significance, and p-values are indicated. **(D)** Western blot analysis of NFAT2 in WT and FKBP2 KO cells. Quantification normalized to GAPDH is shown below. Each dot represents one independent biological replicate (N = 3). Statistical significance was assessed using a paired Student’s t-test. **(E)** Representative high-resolution SIM images of SC-β cells show the localization of NFAT2 (red) and DAPI (blue) in FKBP2 KO and WT cells. Scale bar, 10 µm. **(F)** Immunofluorescence staining of NFAT2 (green) in WT, KO, and KO cells treated with HDAC inhibitor (HDACi, TMP195, 1 µM). DAPI marks nuclei in blue. Scale bar, 100 µm. **(G)** Quantification of normalized NFAT2 mean fluorescent intensity (MFI) under WT, KO, and HDACi-treated KO cells. N = 3 biological repeats. Each dot represents one image; horizontal lines indicate mean ± SD. An unpaired t-test was used to assess significance, and p-values are indicated. **(H)** The schematic summarizes the proposed mechanism. FKBP2 deficiency triggers ER stress. This stress causes an increase in intracellular calcium (Ca²⁺) levels, which activates calcineurin, a calcium and calmodulin-dependent phosphatase. Calcineurin dephosphorylates NFAT2, allowing its translocation to the nucleus, where it can regulate the expression of differentiation-related genes, leading to increased HDAC9 expression, which we hypothesize disrupts the transcription of genes critical for proper β cell development, contributing to increased SC-α cell formation. **(I)** Workflow to generate inducible FKBP2 KO in hPSCs (iKO): iCas9 hPSCs (HUES8) were transfected with a plasmid carrying the piggyBac system and FKBP2-specific sgRNAs. Cells were differentiated to the early SC-β stage (day 21), where Cas9 expression was induced by doxycycline (DOX). Six days post-doxycycline treatment FKBP2 KO was confirmed by immunofluorescence. iKO SC-β cells were analyzed alongside WT, WT + DOX, and iKO without DOX controls. **(J)** Immunofluorescence of FKBP2 (green) in WT, WT+ DOX, iKO, and iKO + DOX-treated cells. DAPI marks nuclei in blue. Scale bar, 100 µm. In the right violin plots for FKBP2 staining for WT, WT + DOX, iKO, iKO + DOX. **(K)** Quantification of FKBP2 staining as shown in panel J. The violin plot displays the distribution of mean fluorescent intensity (MFI) normalized to DAPI, with the median indicated as a dashed line and IQR (25th–75th percentiles) as a dotted line. N = 3 independent experiments is shown. P-values were calculated using an unpaired Student t-test. **(L)** Immunofluorescence of ATF6 (green), NFAT2 (yellow), C-PEP (green), and GCG (red) in WT and iKO + DOX cells. DAPI marks nuclei in blue. Scale bar, 100 µm. **(M)** Violin plots for FKBP2, ATF6, NFAT2, C-PEP, and GCG staining as shown in panel L. The violin plot displays the distribution of mean fluorescence intensity (MFI) normalized to DAPI. The violin plots show the data distribution with the median indicated as a dashed line and IQR (25th– 75th percentiles) as a dotted line. N = 3 independent experiments is shown. P-values were calculated using an unpaired Student t-test.

The NFATs are activated by intracellular calcium ion oscillations and Ca-dependent, calmodulin activates calcineurin, which in turn dephosphorylates NFATs enabling their translocation to the nucleus ^33^. We observed elevated NFAT2 protein expression in early and late KO SC-β cells along with enhanced NFAT2 nuclear localization (**Fig. 7C-E**). The nuclear NFAT2 accumulation, likely induced by Ca^2+^ influx following enhanced ER stress in KO cells, suggests NFAT2 transcriptional activity. Finally, HDAC inhibition in KO cells did not alter NFAT2 protein levels, suggesting that NFAT2 acts upstream of HDAC9 (**Fig 7F** and **G**). Together, these findings point to a functional NFAT2–HDAC9 axis involved in blocking β cell differentiation under FKBP2 deficiency.

In summary, the deletion of FKBP2 induces ER stress which involves the activation of the stress-related proteins PERK and ATF6. This stress response also affects the expression of several ER components, including HSPA5, PPIB, HSP90B1, and ERO1B. The increased ER stress and loss of ER chaperone, FKBP2, collectively cause the intracellular Ca^2+^ level increase. This elevated calcium, in conjunction with calmodulin, activates the enzyme calcineurin. Activated calcineurin dephosphorylates the transcription factor NFAT2, on multiple serine residues causing a conformational change in the NFAT protein. The nuclear localization signal is now exposed, and the nuclear export signal is masked, allowing the dephosphorylated NFAT2 translocation to the nucleus, where it promotes the transcription of β cell associated genes. The increased expression of these genes, including HDAC9, results in preferential α cell formation (**Fig. 7H**).

Lastly, to separate the dual roles of FKBP2 in proinsulin processing and endocrine cell differentiation, we derive doxycycline-inducible FKBP2 KO hPSCs, termed iKO (**Fig. 7I)**. To this end, using piggBac transposase, we integrated sgRNAs targeting exons 1, 2 and 6 of FKBP2 into the genome of iCas9-hUES hPSCs **(Fig. S7A)**. We then differentiated iKO hPSCs to early SC-β cells and induced the Cas9 expression by doxycycline treatment. Six days later, we observed the decreased FKBP2 expression by 42 % (**Fig. 7I-K**). Upon FKBP2 downregulation, iKO SC-β cells exhibited a significant increase in ATF6, NFAT2, C-PEP, and GCG compared to WT controls **(Fig. 7L** and **M**), which points to enhanced ER stress signaling and a shift in endocrine lineage allocation. In contrast, the expression of PDX1, CHGA, INSM1, and HDAC9 remained unchanged **(Fig. S7B** and **C)**, suggesting that FKBP2 loss selectively affects stress responses and β/α cell markers without altering early pancreatic transcription factors or chromatin regulators.

## Discussion

The precise regulation of insulin secretion is fundamentally dependent upon both β cell function and β cell mass. The function of β cells is a multifaceted physiological parameter, encompassing cellular maturation, insulin biosynthesis, glucose sensing, and the secretory process itself. Concurrently, the total β cell mass is determined by developmental accrual and the dynamic equilibrium between β cell proliferation and apoptosis. Even subtle perturbations in insulin synthesis or β cell maturation can predispose individuals to prediabetes; when exacerbated by additional defects, such as insulin resistance, these alterations frequently precipitate the development of overt diabetes.

In this study, we present novel evidence demonstrating that FKBP2, an ER chaperone, not only regulates proinsulin folding, but also, unexpectedly, exerts control over the fate of human pancreatic endocrine cells *in vitro*. Our findings reveal that the absence of FKBP2 significantly disrupts SC-β cell formation, alters the critical α/β cell ratio, and leads to aberrant regulation of the transcription factor NFAT2 and the epigenetic modulator HDAC9. Consequently, these disruptions result in a loss of SC-β cell identity and impaired differentiation.

HDAC9 emerges as a particularly important factor, since its upregulation coincides with conditions of cellular stress and correlates with an increase in glucagon-expressing cells. Our data therefore indicate that HDAC9 is not merely passively associated with lineage imbalance but directly contributes to the skewing of endocrine fate decisions toward the α cell lineage. Moreover, the fact that both MC1568 and TMP195 — two distinct classes IIa inhibitors — converge on similar outcomes highlights the robustness of this mechanism and underscores the functional importance of HDAC9 in governing the equilibrium between α and β cell populations.

Our observations expand upon previous investigations that highlight the indispensable role of ER chaperones, such as HSP90B1, HSP70, GRP78/BiP, PDI, PPIB, in maintaining β cell homeostasis ^34^. Consistent with these earlier findings, which demonstrate that loss-of-function of such chaperones can compromise β cell-specific phenotypic traits ^35,36^, our data further underscore the critical involvement of specific ER chaperones in preserving β cell integrity.

The broader significance of protein folding and ER homeostasis in diabetes is well-established. For instance, misfolding proinsulin mutations, exemplified by the Akita variant, are a well-documented cause of diabetes in both murine models and humans. This pathology arises from a dominant-negative proteotoxicity that impairs insulin production, even when only a subset of insulin alleles is affected, as observed with a single affected allele in mice. Similarly, Wolfram syndrome, a rare monogenic disorder characterized by syndromic diabetes, is caused by pathogenic mutations in the ER-localized WFS1 protein. These genetic predispositions are further corroborated by evidence of activated ER stress response pathways in pancreatic islets from individuals with both type 1 diabetes and type 2 diabetes, collectively emphasizing the pervasive involvement of ER stress in the complex pathogenesis of diverse diabetic conditions.

Deletion of the FKBP2 gene in differentiating SC-β cells resulted in a significant decrease in insulin levels and altered insulin granule structure, without significantly affecting proinsulin levels. This suggests a defect in protein folding and processing rather than a problem with insulin synthesis. This finding is similar to Mutant insulin gene-induced diabetes of youth (MIDY), a form of monogenic diabetes caused by mutations in the insulin gene ^37^. Under normal conditions, proinsulin is the most abundant protein in the ER of β cells, making up about 10% of total protein synthesis under basal conditions and even more under high glucose stimulation. Proinsulin is prone to misfolding, and an unfavorable ER environment can exacerbate this misfolding, contributing to the development and progression of diabetes.

In MIDY, insulin gene mutations cause misfolded proinsulin to accumulate in the ER, leading to ER stress and β cell dysfunction. These mutations are the second most common genetic cause of permanent neonatal diabetes. Of the 70 known diabetes-associated insulin gene mutations, most impair the oxidative folding of proinsulin, preventing its export from the ER. The misfolded mutant proinsulin can also interact with and impair the trafficking of co-expressed WT proinsulin, further reducing insulin production. The severity of proinsulin misfolding and its impact on co-expressed wild-type proinsulin are key factors driving insulin-deficient diabetes.

Finally, protein misfolding and increased ER stress extend far beyond pancreatic β cells. These mechanisms are common threads in the development of numerous other pathologies and insights gained from studying them could pave the way for novel therapeutic strategies for a wide array of diseases, not just diabetes.

## Material and methods

### Cell culture

All experiments with hPSCs were performed in accordance with relevant guidelines and regulations of the Local Bioethics Committee of Medical University in Poznan, Poland. hPSCs with doxycycline-inducible Cas9 expression (HUES8-iCas9) were established in Dr. Huangfu laboratory (MSCK, NYC, USA), through targeting into AAVS1 locus, the inducible Cas9 cassette using TALEN approach ^38^. HUES8-iCas9 hPSCs were cultured in StemFlex (Thermo Fisher Scientific, Netherlands) medium on Geltrex-coated (Thermo Fisher Scientific, Netherlands) plates at 37 °C, under an atmosphere containing 5% CO_2_. Cells were passaged every 3–4 days using PBS-EDTA. Cell lines were routinely tested for mycoplasma with PCR assay and found negative.

### FKBP2 gene deletion in hPSCs

sgRNAs targeting the sequence of exon 1, 2 and 6 for FKBP2 were designed using the Benchling software. The sgRNA sequences are listed in **Table 1**. The sgRNAs were produced in-house using a previously published protocol ^27,39^. Briefly, the PCR template constituted a 120-nucleotide single-stranded DNA including a T7 promoter, sgRNA target-specific, and constant sgRNA sequences. The PCR product was used for *in vitro* transcription to generate sgRNAs for lipofection.

**Table 1.**
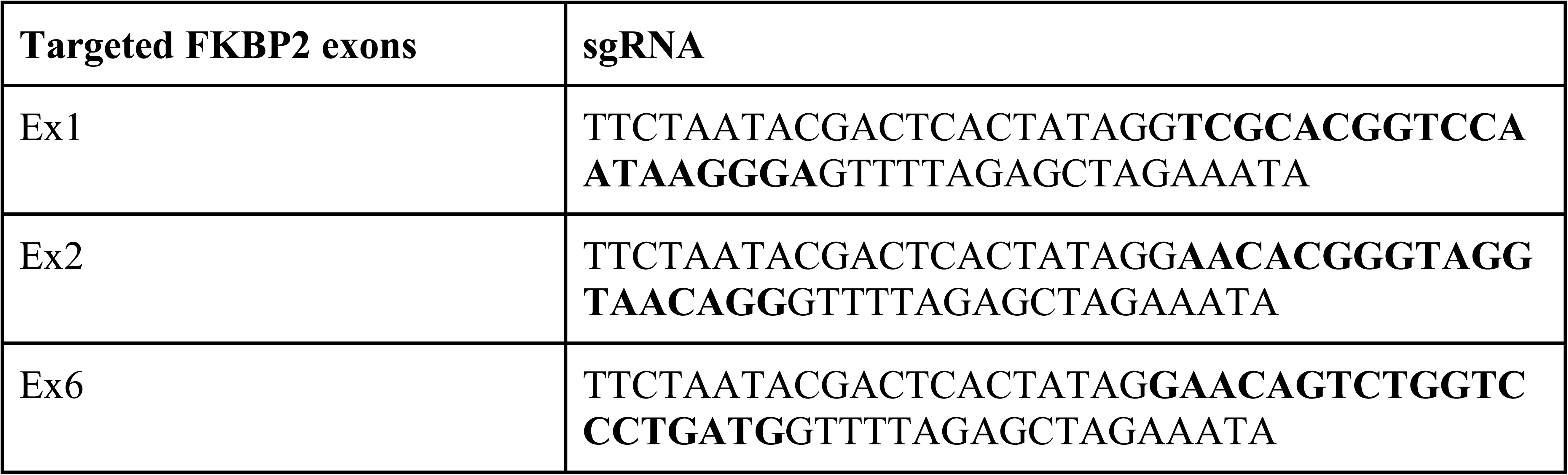
sgRNA sequences used for FKBP2 gene targeting in hPSCs.

Doxycycline-treated HUES8-iCas9 cells were dissociated into single cells using TrypLE Select (Thermo Fisher Scientific, Netherlands), and seeded in E8 medium (Thermo Fisher Scientific, Netherlands) with 10 µM Y-27632 (Peprotech, USA) into Geltrex-coated 24-well plate at the density of 1.5 × 10⁵ cells/well. The cells were reversely transfected using Lipofectamine RNAiMax (Thermo Fisher Scientific, Netherlands) and cultured for 2 days. Subsequently, the cells were plated at single-cell density onto Geltrex-coated 6 mm dishes, cultured for the following 7 days, and single colonies were transferred into a 96-well plate. The PCR was performed on genomic DNA isolated from individual colonies with the primers flanking the *FKBP2* gene sequence. Homozygous clones with desired mutations were selected based on amplified DNA electrophoresis and Sanger sequencing.

### hPSC pancreatic differentiation

For *in vitro* pancreatic differentiation, we used a previously established protocol with modifications. hPSCs were dispersed into single-cell suspensions with TrypLE Select (Thermo Fisher Scientific, Netherlands) and plated at a density of 3.5 × 10^5^ cells/cm² in E8 (Thermo Fisher Scientific, Netherlands) medium supplemented with 10 µM Y-27632 (Peprotech, USA) on low-adhesion non-treated 6-well plates (Eppendorf, Germany) on an orbital shaker at 37 °C, 110 rpm, and under an atmosphere containing 5% CO₂. On subsequent days, the medium was changed to fresh E8 medium without Y-27632. The next day the spheres were washed in DMEM/F12 medium (Corning, USA) and the medium was changed, as follows:

#### Basal medium

**S1 –** MCDB131 (Corning, USA), 1% Glutamax (Thermo Fisher Scientific, Netherlands) + 1% Pen/Strep (Thermo Fisher Scientific, Netherlands), 2.44 mM glucose (Merck Millipore, USA), 29 mM sodium bicarbonate (Merck Millipore, USA), 2% BSA (Capricorn Scientific, USA), 2 μl/100 ml ITS-X (Thermo Fisher Scientific, Netherlands).

**S2 –** MCDB131 (Corning, USA), 1% Glutamax (Thermo Fisher Scientific, Netherlands) + 1% Pen/Strep (Thermo Fisher Scientific, Netherlands), 2.44 mM glucose (Merck Millipore, USA), 13 mM sodium bicarbonate (Merck Millipore, USA), 2% BSA (Capricorn Scientific, USA), 2 μl/100 ml ITS-X (Thermo Fisher Scientific, Netherlands).

**S3 –** MCDB131 (Corning, USA), 1% Glutamax (Thermo Fisher Scientific, Netherlands) + 1% Pen/Strep (Thermo Fisher Scientific, Netherlands), 2.44 mM glucose (Merck Millipore, USA), 13 mM sodium bicarbonate (Merck Millipore, USA), 2% BSA (Capricorn Scientific, USA), 500 μl/100 ml ITS-X (Thermo Fisher Scientific, Netherlands).

**S4** – MCDB131 (Corning, USA), 1% Glutamax (Thermo Fisher Scientific, Netherlands) + 1% Pen/Strep (Thermo Fisher Scientific, Netherlands), 2.44 mM glucose (Merck Millipore, USA), 29 mM sodium bicarbonate (Merck Millipore, USA), 2% BSA (Capricorn Scientific, USA), 500 μl/100 ml ITS-X (Thermo Fisher Scientific, Netherlands), 5 µg/ml heparin sulfate (Sigma Aldrich, USA),

**S5** – MCDB131 (Corning, USA), 1% Glutamax (Thermo Fisher Scientific, Netherlands) + 1% Pen/Strep (Thermo Fisher Scientific, Netherlands), 2.44 mM glucose (Merck Millipore, USA), 2% BSA (Capricorn Scientific, USA), 260 nM zinc sulfate (Merck Millipore, USA), 5 µg/mL heparin sulfate (Sigma Aldrich, USA), 1x Trace Elements A (Corning, USA), 1x Trace Elements B (Corning, USA).

**Day 1:** S1 + 3 µM CHIR99021 (Peprotech, USA) + 100 ng/ml Activin A (Peprotech, USA) + 250 μM vitamin C (Sigma Aldrich, USA).

**Days 2–3:** S1 + 100 ng/ml Activin A + 250 μM vitamin C.

**Days 4–6:** S2 + 50 ng/ml KGF (Peprotech, USA) + 1.25 μM IWP2 (Selleckchem, USA) + 250 μM vitamin C.

**Day 7:** S3 + 50 ng/ml KGF + 2 μM retinoid acid (Peprotech, USA) + 500 nM PdBu (Tocris, United Kingdom) + 250 nM SANT-1 (Tocris, United Kingdom) + 200 nM LDN193189 (Peprotech, USA) + 250 μM vitamin C + 10 μM Y-27632.

**Days 8-12:** S3 + 5 ng/ml Activin A + 50 ng/ml KGF + 100 nM retinoid acid + 250 nM SANT-1 + 1.25 μM IWP2 + 100 ng/ml EGF (Peprotech, USA) + 10 mM nicotinamide (Sigma Aldrich, USA) + 250 μM vitamin C.

**Day 13-19:** S4 + 100 nM retinoic acid (Peprotech, USA) + 250 nM SANT-1 (Tocris, United Kingdom) + 1 µM LY-411575 (Adooq Bioscience, USA) + 1 µM T3 (Sigma Aldrich, USA) + 10 µM Alk5iII (Adooq Bioscience, USA) + 20 ng/mL betacellulin (Peprotech, USA) + 250 μM vitamin C.

**Day 20 and beyond:** S5 without additional components.

### Western blot analysis

Cells were lysed in the buffer composed of 60 mM Tris (Bio-Shop, Canada), 2% SDS (Bio-Shop, Canada), 10% sucrose (Bio-Shop, Canada), and 1% protease inhibitor cocktail (Sigma Aldrich, USA), followed by sonication for 15 seconds in 8 cycles with 1 min pause between cycles and centrifugation at 12,000 g for 10 min. Protein concentration was determined using a NanoDrop spectrophotometer. 30 µg of cell lysates in Bolt LDS buffer (Thermo Fisher Scientific, Netherlands) were incubated for 5 min at 95 °C and then loaded on a 12% SDS-PAGE gel. Electrophoresis was carried out at 75 V for the first 15 min and 125 V for an additional 1h, with a constant current intensity of 30 mA. Proteins were transferred onto the PVDF (Thermo Fisher Scientific, Netherlands) membrane using the BioRad Trans-Blot Turbo Transfer System for 10 min. Membranes were blocked with 3% BSA in TBS-Tween 20 followed by overnight incubation with a primary antibody. On the next day, membranes were probed with HRP-conjugated secondary antibodies. All antibodies are listed in **Table 2**. Western blot results were visualized using the enhanced chemiluminescent visualization (ECL) system (Thermo Scientific Pierce) or SuperSignal West Femto Maximum Sensitivity Substrate (Thermo Fisher Scientific, Netherlands). Signal detection was conducted using the G: Box System (Syngene, India). Western blots were quantified in ImageJ software.

**Table 2.**
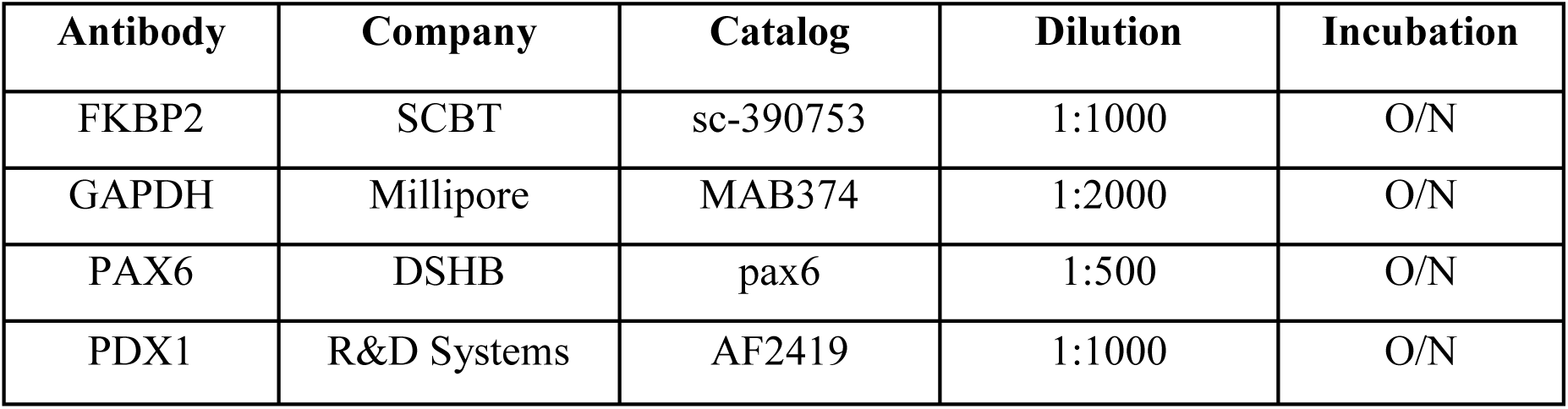
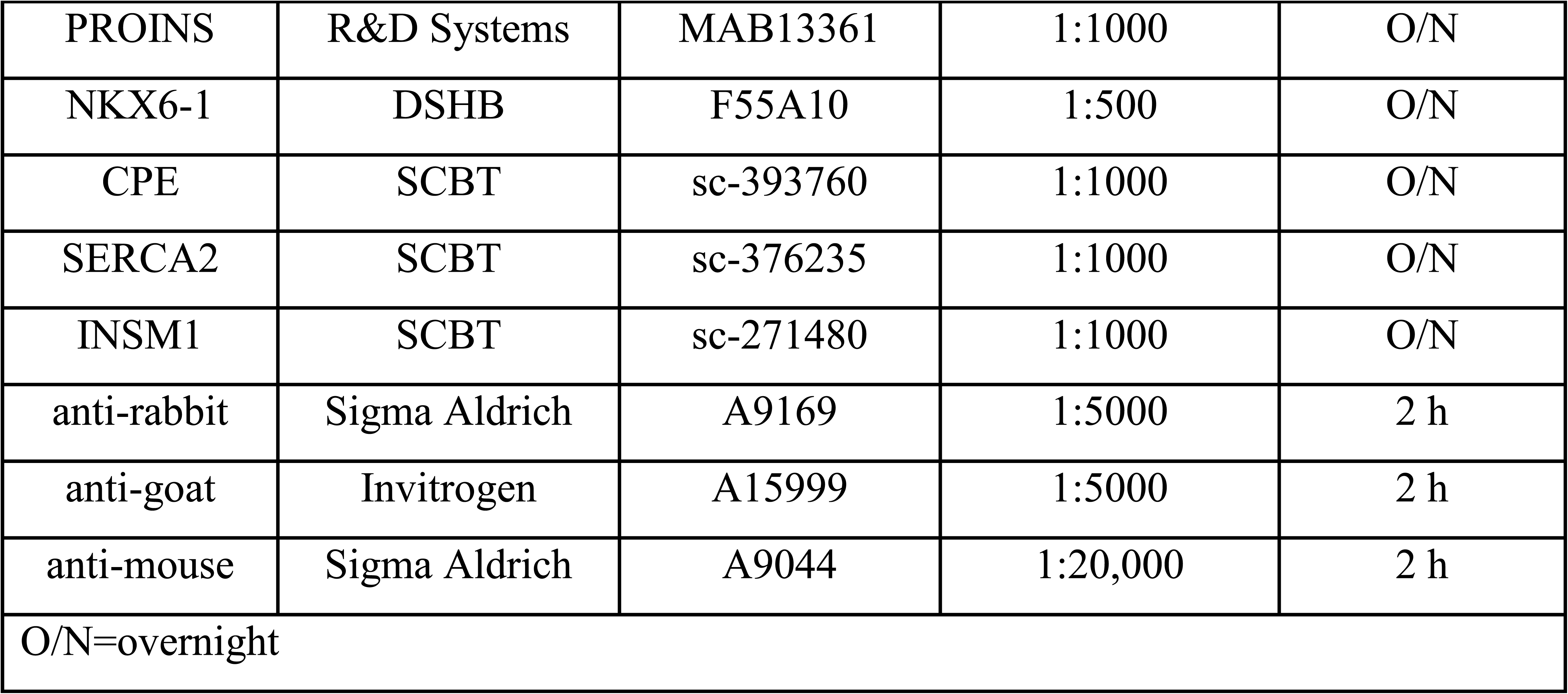
Antibodies used for western blot analysis.

**Cytosolic calcium level** was determined using calcium Fluo-4 AM dye (Thermo Fisher Scientific, Netherlands). Cells were stained with 2.5 µM concentration of Fluo-4 AM and incubated at 37°C for 30 min in the dark. Immediately after staining, cells were washed twice with DPBS and analyzed with 488 nm excitation using a Leica Stellaris 8 confocal microscope. Data were analyzed using the Leica LAS X.

### Single-cell RNA sequencing

WT and KO spheres at the early (day 21) and late (day 31) β cell stage, were dissociated into single cells using 1% trypsin in PBS at 37 °C. Libraries for scRNA-seq were conducted with 10x Genomics kit according to the manufacturer’s protocol. The single-cell transcriptome was marked with UMI barcode using a droplet-based method in Chromium (10x Genomics, USA). For quality and quantity assessment of cDNA libraries, Agilent TapeStation was used with High Sensitivity D1000 ScreenTape (Agilent Technologies, USA). Libraries were paired-end sequenced with a depth of 40,000 reads per cell. 10,355 WT cells and 6419 KO cells, as well as 5,200 WT and 5,800 KO cells were sequenced for WT and KO the early and late SC-β cell stages, respectively. Bioinformatic analyses were conducted with scripts for CellRanger (10x Genomics, USA), including data aggregation with the “cellranger agrr” algorithm. The scRNA-seq data have been deposited in the NCBI GEO database under accession number GSE304060. Single-cell data clustering and visualization with the UMAP algorithm, analysis of specific markers or DEGs for each cluster (using Wilcoxon Rank Sum test), and violin and feature plots with gene expression patterns within the clusters were performed in R Studio using Seurat Package v4.3.0. Enriched pathways and processes among markers for each cluster and DEGs were identified via the KEGG, GO, Biological Processes (BP), or Wiki Pathways databases. The functional enrichment analysis was performed in the GeneCodis interactive platform. A pseudotime trajectory was performed using Monocle3. The significance cutoff value (adj p-value < 0.05) was utilized in functional enrichment analysis.

### Immunofluorescence staining

Human pancreas sections were processed at Baylor College of Medicine, Houston, TX (USA) with IRB H-3097 approval granted to Dr. Malgorzata Borowiak. Donor identities were encrypted, and the data were analyzed anonymously. The human 13-week fetal pancreas samples were fixed in 4% paraformaldehyde/PBS for 4 h, washed with PBS, soaked in 30% sucrose, and embedded in TissueTek. Sections (12 μm-thick) were cut onto Superfrost Plus-coated glass slides and stored at −80 °C.

Cell monolayers were fixed with 4% paraformaldehyde/PBS for 15 min at room temperature and washed with PBS. Fixed cells were permeabilized by 0.5% Triton X-100 (BioShop, Canada) in PBS for 15 min, blocked for 45 min in 3% BSA (BioShop, Canada) + 0.1% Tween 20 (BioShop, Canada) in PBS and then incubated overnight at 4 °C with primary antibodies diluted in 5% normal donkey serum (NDS, Jackson Immunoresearch, UK). The next day, cells were washed three times in 0.1% Tween 20 in PBS and incubated with secondary antibodies conjugated with Alexa Fluor 488, TRITC, or Alexa Fluor 647 (diluted 1:400 in 5% NDS) for 2 h at room temperature. The excess secondary antibody was removed by two washes in 0.1% Tween 20 in PBS, and the samples were incubated with DAPI (Sigma Aldrich, USA) as a counterstain. To stain lipids, 1 µg/mL Nile Red (AAT Bioquest, USA) was added to the medium, and cells were incubated at 37 °C for 15 minutes, then washed with PBS and fixed.

3D Spheres were fixed with 4% paraformaldehyde/PBS for 45 min at 4 °C, washed once with 0.1% Tween 20 in PBS, and suspended in spheroid wash buffer (SWB: 2% BSA, 0.1% Triton X-100, 0.025% SDS, 1 × PBS). Fixed spheres were blocked in SWB for 45 min at 4 °C and incubated overnight with a primary antibody at 4 °C on an orbital shaker. The next day, the spheres were washed 3 times with SWB solution for 1 h each at 4 °C and incubated with secondary antibodies conjugated with Alexa Fluor 488, TRITC, or Alexa Fluor 647 diluted 1:400 and DAPI (1:10,000) in SWB, overnight at 4 °C on orbital shaker. On the next day, the excess of the secondary antibody was removed by 2 washes in 0.1% Tween 20 in PBS, first rapid, second for 1 h at 4 °C on an orbital shaker. For imaging, spheres were kept in PBS on a high-content imaging glass-bottom 96-well plate (Corning, USA). The primary and secondary antibodies used in the study are listed in **Table 3**.

**Table 3.** Antibodies used for immunofluorescence staining.

| Antigen | Company | Catalog | Dilution | Primary/<br>Secondary | Host | Reactivity |
| --- | --- | --- | --- | --- | --- | --- |
| ATF6 $\alpha$ | SCBT | sc-166659 | 1:100 | Primary | Mouse | Hu, Mus, Rat |
| CHGA | Abcam | ab15160 | 1:200 | Primary | Rabbit | Hu |
| CPE | SCBT | sc-393760 | 1:100 | Primary | Mouse | Hu, Mus, Rat |
| C-PEP | DSHB | GN-ID4 | 1:40 | Primary | Rat | Hu, Monkey |
| FKBP2 | SCBT | sc-390753 | 1:100 | Primary | Mouse | Hu, Mus, Rat |
| FOXA2 | R&D Systems | AF2400 | 1:100 | Primary | Goat | Hu |
| GCG | SCBT | sc-514592 | 1:100 | Primary | Mouse | Hu, Mus, Rat |
| HDAC9 | Abcam | AB109446 | 1:400 | Primary | Rabbit | Hu |
| INS | Cell signaling | 3014 | 1:100 | Primary | Rabbit | Hu, Mus, Rat |
| INS | SCBT | sc-8033 | 1:100 | Primary | Mouse | Hu, Mus, Rat |
| INSM1 | SCBT | sc-271480 | 1:100 | Primary | Mouse | Hu, Mus, Rat |
| NFAT2 | Abcam | AB25916 | 1:300 | Primary | Rabbit | Hu |
| NKX6-1 | DSHB | F55A10 | 1:20 | Primary | Mouse | Hu, Mus |
| OCT3/4 | SCBT | sc-5279 | 1:100 | Primary | Mouse | Hu, Mus, Rat |
| P115 | SCBT | sc-48363 | 1:100 | Primary | Mouse | Hu, Mus, Rat |
| PC1/3 | Abcam | AB220363 | 1:100 | Primary | Rabbit | Hu, Mus, Rat |
| PDX1 | R&D Systems | AF2419 | 1:100 | Primary | Goat | Hu |
| PERK | SCBT | sc-377400 | 1:100 | Primary | Mouse | Hu, Mus, Rat |
| PROINS | R&D Systems | MAB13361 | 1:100 | Primary | Mouse | Hu, Mus |
| SERCA2 | SCBT | sc-376235 | 1:50 | Primary | Mouse | Hu, Mus, Rat |
| SST | SCBT | sc-7819 | 1:100 | Primary | Goat | Hu |
| VIM | DSHB | H5 | 1:100 | Primary | Mouse | Hu, Mus, Rat,<br>Chick |
| AlexaFluor<br>488 | Jackson<br>Immunoresearch | 711-545-152 | 1:400 | Secondary | Donkey | Anti-Rabbit IgG<br>(H+L) |
| AlexaFluor<br>488 | Jackson<br>Immunoresearch | 715-545-147 | 1:400 | Secondary | Donkey | anti-Goat IgG<br>(H+L) |
| AlexaFluor<br>488 | Jackson<br>Immunoresearch | 705-545-150 | 1:400 | Secondary | Donkey | anti-Mouse IgG<br>(H+L) |
| AlexaFluor<br>647 | Jackson<br>Immunoresearch | 705-605-147 | 1:400 | Secondary | Donkey | anti-Goat IgG<br>(H+L) |
| AlexaFluor<br>647 | Jackson<br>Immunoresearch | 711-605-152 | 1:400 | Secondary | Donkey | anti-Rabbit IgG<br>(H+L) |
| AlexaFluor<br>647 | Jackson<br>Immunoresearch | 715-605-151 | 1:400 | Secondary | Donkey | anti-Mouse IgG<br>(H+L) |
| TRITC | Jackson<br>Immunoresearch | 711-025-152 | 1:400 | Secondary | Donkey | anti-Rabbit IgG<br>(H+L) |
| TRITC | Jackson<br>Immunoresearch | 715-025-150 | 1:400 | Secondary | Donkey | anti-Mouse IgG<br>(H+L) |
| TRITC | Jackson<br>Immunoresearch | 705-025-147 | 1:400 | Secondary | Donkey | anti-Goat IgG<br>(H+L) |

### Microscopy imaging

Images were obtained with an epifluorescence, confocal, super-resolution, or light sheet microscope. Bright-field images were taken with a Leica DM IL-Led (Leica, Germany) microscope with N Plan Fluor 4×/0.12, N Plan Fluor 10×/0.30, N Plan Fluor 20×/0.40 and N Plan Fluor 40×/0.60 lenses and a JENOPTIK GRYPHAX series ProgRes camera (JENOPTIK, Germany). Fluorescence images were obtained with a confocal microscope Nikon A1Rsi (Nikon, Germany) microscope with Plan Fluor 4×/0.13, Plan Apo 10×/0.45 DIC N1, Plan Apo VC 20×/0.75 DIC N2, Apo 40×/1.25 WI λS DIC N2, and Plan Apo VC 60×/1.4 Oil DIC N2 lenses and with Nikon NIS Elements AR 5.21.01 64-bit software (Nikon, Germany) and Leica Stellaris 8 confocal microscope (Leica Microsystems, Germany) equipped with white light laser (WLL) excitation and acousto-optical tunable filter (AOTF) detection. Images were acquired using the following objectives: HC PL APO 10×/0.40 CS2, HC PL APO 20×/0.75 IMM CORR CS2, HC PL APO 40×/1.30 OIL CS2, and HC PL APO 63×/1.40 OIL CS2. The system was operated with Leica LAS X (v.4.3 or higher) software in 64-bit configuration. Image acquisition included hybrid detector (HyD S) modules, enabling spectral detection and high sensitivity. For z-stacks and time-lapse imaging, acquisition settings were optimized to maintain signal-to-noise ratio and avoid photobleaching. Super-resolution microscopy was performed with a ZEISS Elyra 7 with Lattice SIM (Zeiss, Germany) microscope with Plan-Apochromat 40x/1.4 Oil DIC M27 (Immersol 518F – 23 °C/30 °C) and Plan-Apochromat 63×/1.40 Oil DIC f/ELYRA (Immersol 518F – 23 °C/30 °C) lenses and with ZEN Black 3.1 software (for imaging) and ZEN Blue 3.1 software for data analysis (Zeiss, Germany) or with light sheet fluorescence microscopy (LSFM) ZEISS Lightsheet 7 (Zeiss, Germany) with W Plan-APO 20x/1,0 DIC lens, in water chamber and with ZEN Black 3.1 software for imaging and ZEN Blue 3.1 software for data analysis (Zeiss, Germany).

### Transmission electron microscopy

To analyze the granular ultrastructure, WT and KO SC-β cells were fixed at room temperature for 2 h in a mixture containing 1.25% paraformaldehyde, 2.5% glutaraldehyde, and 0.03% picric acid in 0.1 M sodium cacodylate buffer (pH 7.4). The samples were then washed in 0.1 M cacodylate buffer and postfixed at room temperature with a mixture of 1% OsO_4_/1.5% KFeCN_6_ once for 2 h and then once for 1 h. After washing with water, the samples were stained in 1% aqueous uranyl acetate for 1 h, washed, and subsequently dehydrated. A 1 h incubation in propylene oxide was followed by infiltration overnight in a 1:1 mixture of propylene oxide and TAAB Epon, after which the samples were embedded in TAAB Epon. The cut sections were then stained with 0.2% lead citrate. A JEOL 1200EX transmission electron microscope or a TecnaiG2 Spirit BioTWIN was used to analyze the samples.

### ER stress induction

Cells were cultured under standard conditions at 37 °C in an incubator with 5% CO₂. ER stress was induced by treating cells with DTT (MedChemExpress, USA), a reducing agent that disrupts disulfide bond formation, leading to the accumulation of misfolded proteins in the ER and activation of the UPR. WT and KO cells were treated with 1 mM DTT at 37 °C for 8 days, followed by two washes with PBS and immunofluorescence staining.

### HDAC9 inhibition

Cells were cultured under standard conditions at 37 °C in an incubator with 5 % CO₂. HDAC9 inhibition was performed only in KO cells by treatment with TMP195, a selective class IIa histone deacetylase (HDAC) inhibitor targeting HDAC4, HDAC5, HDAC7, and HDAC9, with the highest potency toward HDAC9. KO cells were treated with 1 µM TMP195 at 37 °C for 6 days, followed by two washes with PBS and immunofluorescence staining. WT untreated and KO untreated cells were used as controls.

### Live-cell imaging and analysis

The IncuCyte live-cell imager (Sartorius, Germany) was used to track living cells. hPSCs were seeded at cell density 1.5 × 10^4^/cm^2^ on 24-well Geltrex-coated plates and cultured in IncuCyte for a maximum of 6 days at 37 °C under an atmosphere containing 5% CO_2_. Photomicrographs were taken every 2 h and the confluence was measured in real time with the IncuCyte Base Analysis Software (Sartorius, Germany). Cell growth was monitored by analyzing the change in the area of an image occupied by the cells (confluency in %) over time.

### Flow cytometry

Cells were dissociated with TrypLE Select (Thermo Fisher Scientific, Netherlands) to obtain a single-cell suspension and fixed by incubation with 4% paraformaldehyde in PBS with 0.1% saponin for 45 min at 4 °C and washed with 0.1% saponin/1% BSA in PBS (SBP). The resuspended cells were incubated overnight at 4 °C on the roller with primary antibodies diluted in SBP. The following day, the cells were washed twice with SBP and incubated with secondary antibodies conjugated with TRITC, Alexa Fluor 488, or Alexa Fluor 647 (Jackson ImmunoResearch, UK) diluted between 1:400 and 1:600 in SBP for 1 h at room temperature. The excess secondary antibodies were removed by washing once with PBS and the cells, resuspended in PBS, were used directly for flow cytometry analysis. Flow cytometry data were acquired with a CytoFLEX Flow (Beckman Coulter, USA), or NovoCyte Flow Cytometer (Agilent Technologies, USA). Flow cytometry data analysis, including gating, quantification, and the generation of density plots/histograms, was performed with FlowJo Software v10 (BD, USA) or NovoExpress Software v1.6.2 (Agilent Technologies, USA). The primary and secondary antibodies used in the study are listed in **Table 4**.

**Table 4.** Antibodies used for flow cytometry.

| Antigen | Company | Catalog | Dilution | Primary/<br>Secondary | Host | Reactivity |
| --- | --- | --- | --- | --- | --- | --- |
| ETV1 | Invitrogen | PA5-67447 | 1:300 | Primary | Rabbit | Hu |
| INS | Cell signaling | 3014 | 1:300 | Primary | Rabbit | Hu, Mus, Rat |
| NKX6-1 | DSHB | F55A10 | 1:600 | Primary | Mouse | Hu, Mus |
| OCT3/4 | SCBT | sc-5279 | 1:200 | Primary | Mouse | Hu, Mus, Rat |
| NANOG | R&D Systems | AF1997 | 1:200 | Primary | Goat | Hu |
| PDX1 | R&D Systems | AF2419 | 1:1000 | Primary | Goat | Hu |
| AlexaFluor<br>488 | Jackson<br>Immunoresearch | 705-545-150 | 1:5000 | Secondary | Donkey | anti-Mouse IgG<br>(H+L) |
| AlexaFluor<br>647 | Jackson<br>Immunoresearch | 705-605-147 | 1:5000 | Secondary | Donkey | anti-Goat IgG<br>(H+L) |
| AlexaFluor<br>647 | Jackson<br>Immunoresearch | 711-605-152 | 1:5000 | Secondary | Donkey | anti-Rabbit IgG<br>(H+L) |

### Doxycycline-inducible FKBP2 overexpression in hPSCs

*FKBP2* cDNA was amplified from total WT cDNA with primers consisting of *FKBP2* coding sequence, FLAG sequence at the 3’ end, and adapters to the plasmid. Primers consisted of the following sequences:

F: CTTTAAAGGAACCAATTCAGccaccATGAGGCTGAGCTGGTTCCG
R: GCTGGGTCTAGATATCTCGATTACAGCTCAGTTCGTCGCTCTA

Primers were designed using the Benchling Assembly Wizard tool. *FKBP2* cDNA was amplified via PCR using Q5 polymerase (NEB, USA) following the manufacturer’s instructions, with two-step PCR cycling conditions. The cycling conditions were: 98 °C for 30 s, followed by 15 cycles of 98 °C for 15 s, 63 °C for 20 s, 72 °C for 90 s, 25 cycles of 98 °C for 15 s, 72 °C for 15 s, 72°C for 90 s, followed by final extension at 72 °C for 5 min. Monarch DNA Gel Extraction Kit (NEB, USA) was used for the resulting *FKBP2* PCR product purification. Gateway pENTR 1A plasmid (Invitrogen, Netherlands) was digested with the restriction enzymes SalI and XhoI (Thermo Fisher Scientific, Netherlands). To ligate the *FKBP2* PCR product with vector NEBuilder HiFi DNA Assembly Cloning Kit (NEB, USA) was used. The resulting Gateway pENTR 1A plasmid containing the *FKBP2* cDNA sequence was subjected to an LR recombination reaction with plasmid PB-TA-ERP2 ^40^. PB-TA-ERP2 was a gift from Knut Woltjen (Addgene plasmid # 80477 and Addgene plasmid # 80477). WT and KO hPSCs were transfected with the FKBP2-OE plasmid and pCMV-hyPB plasmid ^41^ (gift from Dr. Allan Bradley) carrying transposase enzyme using Lipofectamine 3000 kit (Thermo Fisher Scientific, Netherlands), following manufacturer’s instructions. On the day of transfection, cells were treated with 1 µg/mL puromycin to select positively transformed cells. To induce FKBP2 overexpression, cells were treated with 1 µg/mL doxycycline for 6 days.

### Doxycycline-inducible FKBP2 KO in hPSC-derived β cells

To generate a time-regulated KO of *FKBP2* gene in SC-β cells, we employed a CRISPR/Cas9 system under doxycycline control, utilizing the Tet-On transcriptional activation mechanism and PiggyBac transposon-mediated integration. We used the same sgRNAs targeting the first, second, and sixth exons of the *FKBP2* gene as for generation of global FKBP2 KO (**Table 1**). These sgRNAs, separated by spacer sequences, were synthesized as a single DNA fragment flanked by regions homologous to the PB-gRNA-Puro plasmid (Addgene #121121). The insert was amplified by PCR using forward primer F: ATCTTGTGGAAAGGACGAAA and reverse primer R: GAATACTGCCATTTGTCTCGAG. PCR cycling conditions were initial denaturation at 98 °C for 30 s, followed by 30 cycles of 98 °C for 10 s, 61 °C for 30 s, and 72 °C for 30 s, with a final extension at 72 °C for 2 min. The PB-gRNA-Puro plasmid was digested with BbsI (Thermo Fisher Scientific, Netherlands) and EcoRI (Thermo Fisher Scientific, Netherlands) restriction enzymes in Tango Buffer (Thermo Fisher Scientific, Netherlands) for 4 h at 37 °C, followed by heat inactivation at 65 °C for 20 min. The purified insert and linearized plasmid were assembled using the NEBuilder HiFi DNA Assembly Kit. Competent *E. coli* NEB 10-beta were transformed with the assembled plasmid by heat shock. Transformed cells were recovered in LB medium containing 100 µg/ml ampicillin for 1 h at 37 °C, then plated onto LB agar plates with 100 µg/ml ampicillin and incubated overnight at 37 °C. Individual colonies were screened by colony PCR using the F and R primers listed above. Positive clones were expanded in 50 ml LB + 100 µg/ml ampicillin overnight at 37 °C. Plasmids were purified and quantified on a NanoDrop spectrophotometer. HUES8-iCas9 hPSCs were seeded at 70 % confluency on Geltrex-coated 24-well plates in E8 medium. Transfections were performed with Lipofectamine 3000 according to the manufacturer protocol. To select for stably integrated constructs, 2 µg/ml puromycin (BioShop, Canada) was added for 24 h. Viable puromycin-resistant colonies were genotyped by PCR and expanded in E8 medium. For temporal control of FKBP2 KO during differentiation, 1 µg/ml doxycycline was added to the culture medium on day 21 of differentiation, triggering Tet-On–mediated induction of Cas9 expression and sgRNA-guided editing. KO efficiency was evaluated at the protein level by immunofluorescence microscopy, comparing doxycycline-induced samples with non-induced controls to confirm FKBP2 depletion in SC-β cells.

## Statistics

All graphs were generated using GraphPad Prism v8.4.2 for Windows (GraphPad Software, Boston, Massachusetts, USA). Statistical analyses were conducted using unpaired two-tailed Student’s t-tests or one-way ANOVA for multiple comparisons. Data are presented as means ± SD.

## Author contribution

E. U. - experimental design and execution, including FKBP2 KO hPSC generation, pancreatic differentiation, scRNA-seq analysis of WT and KO SC-β cells, data acquisition and analysis, figure preparation and manuscript writing; W. J. Sz. - data analysis and figure preparation; E. S.- scRNA-seq and pseudotime analysis; A. J. - performed TEM analysis; M. G. - performed flow cytometry, M. Ba - participated in experimental execution, including pancreatic differentiation, derived iKO hPSC lines, M. M. - conceptualization, data analysis, manuscript editing, M. B. - conceptualization, experimental design, data analysis, manuscript writing, and funding acquisition.

## Acknowledgments

We thank Prof. Carmen Birchmeier, and Dr. Thomas Muller MDC, Berlin, Germany for sharing INSM1 antibodies. We also thank Dr. Jolanta Chmielowiec for fetal pancreas sample preparation as well as discussions and comments. We also thank Dr. Knut Woltjen for sharing plasmid PB-TA-ERP2 via ADDGENE (#80477). We also thank Dr. Pablo Navarro for sharing plasmid PB-gRNA-Puro via ADDGENE (#121121). Finally, we would like to thank all members of Borowiak lab for valuable discussions and daily support.

## Data availability

Raw RNA-seq data generated during the study have been deposited in the NCBI GEO database under accession number GSE304060. Raw image files are available from the corresponding author upon a reasonable request. The authors declare that cell lines are available for the research community upon a request from the corresponding author.

