## Supplementary-Figures 1_7-and-Table-Legends for "ER stress and cell fate: FKBP2 regulates proinsulin folding and α- vs. β cell differentiation via NFAT and HDAC9"

### Supplementary information

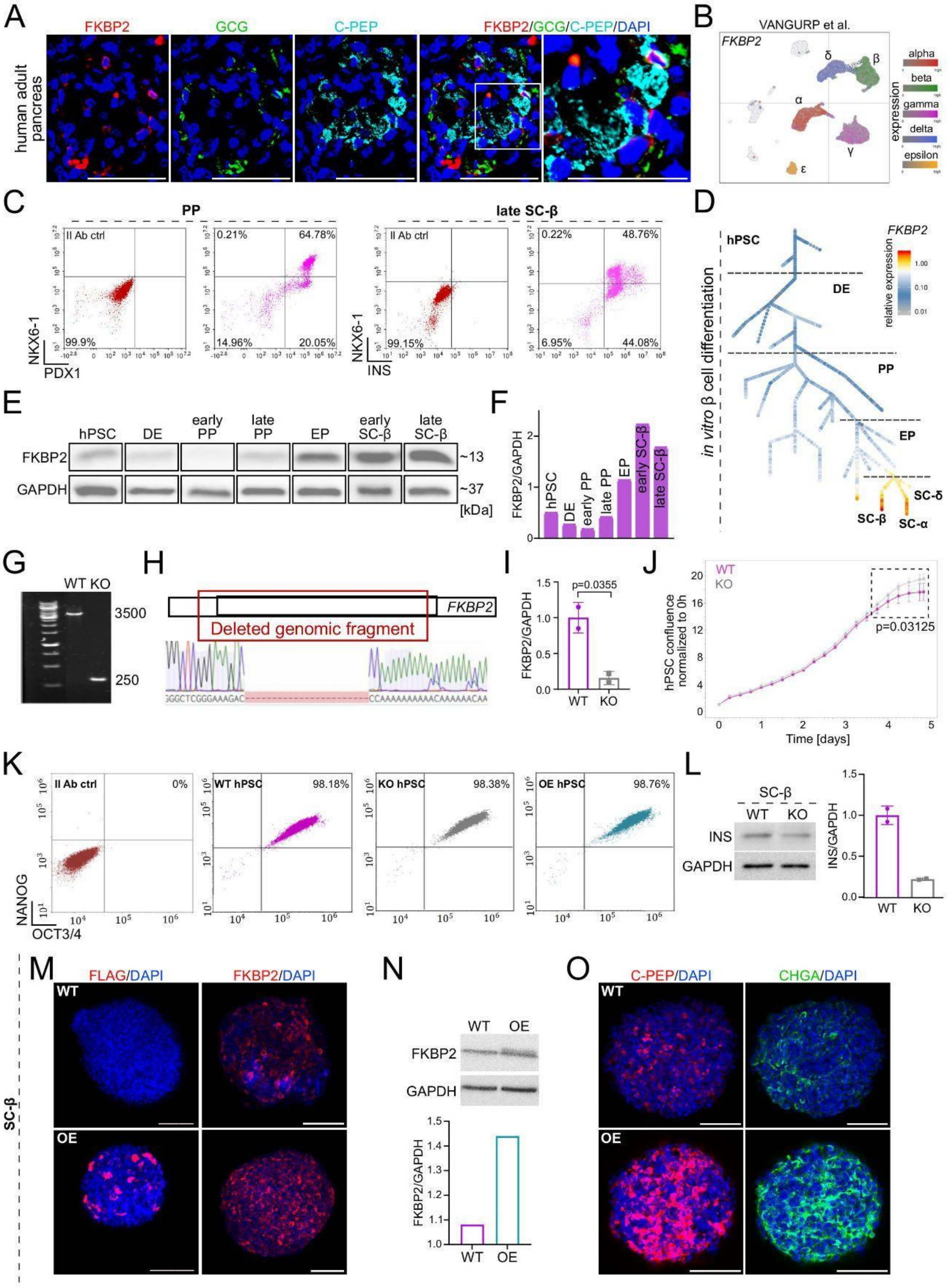

**Supplementary Figure 1: FKBP2 expression dynamics during hPSC pancreatic differentiation and FKBP2 KO and OE hPSC initial assessment**

**(A)** Immunofluorescence staining of human adult pancreas tissue sections shows expression of FKBP2 (red), glucagon (GCG, green), and C-peptide (C-PEP, cyan). DAPI marks nuclei (blue). The boxed region indicates an area of interest with co-localization. Scale bar, 50 µm.

**(B)** Analysis of single-cell RNA-seq (scRNA-seq) data from human adult pancreas showing *FKBP2* mRNA expression across pancreatic cells based on van Gurp *et al.* [9].

**(C)** Representative flow cytometry analysis of PDX1 and NKX6-1 co-expression at PP stage and NKX6-1 and insulin (INS) at SC-β cell stage of WT hPSC pancreatic differentiation.

**(D)** ScRNA-seq data analysis and representation of the relative *FKBP2* mRNA expression during *in vitro* hPSC differentiation to SC-β cells based on Weng *et al*. [13].

**(E)** Western blot analysis of FKBP2 protein expression during WT hPSC differentiation to SC-β cells. GAPDH is shown as a loading control.

**(F)** Quantification of FKBP2 protein levels normalized to GAPDH from western blots.

**(G)** Gel electrophoresis of genomic PCR products confirming the deletion of *FKBP2* exons 1-6 in FKBP2 KO hPSCs.

**(H)** Sanger sequencing results show the deleted *FKBP2* genomic fragment in FKBP2 KO cells.

(**I**) Quantification of FKBP2 protein levels normalized to GAPDH in WT and FKBP2 KO SC-β cells. Each dot represents one biological replicate, N = 2 independent biological replicates; bars show mean ± SD.

**(J)** Quantification of WT and KO hPSC confluency over 5 days of culture. Data represent mean ± SD from N = 4 independent biological replicates. Statistical significance was assessed using a paired two-tailed Student's t-test.

**(K)** Flow cytometry analysis of NANOG and OCT3/4 expression in WT, FKBP2 KO, and FKBP2 OE hPSCs, along with secondary antibody control.

**(L**) Western blot analysis and quantification of insulin (INS) protein expression in WT and FKBP2 KO SC-β cells. GAPDH is shown as a loading control. N=2

(**M**) Immunofluorescence staining for FLAG and FKBP2 protein expression in WT and FKBP2 overexpressing (OE) SC-β cells. DAPI marks nuclei. OE cells were treated with 1 µg/mL doxycycline at day 20 and stained at day 26 of differentiation. Scale bar, 100 µm.

(**N**) Western blot analysis and quantification of FKBP2 protein expression in WT and FKBP2-overexpressing (OE) SC-β cells, following 1 µg/mL doxycycline treatment for 6 days. GAPDH is shown as a loading control.

(**O**) Immunofluorescence staining for C-peptide (C-PEP, red) and chromogranin A (CHGA, green) in WT and FKBP2-overexpressing (OE) SC-β cells, following 1 µg/ml doxycycline treatment for 6 days. Scale bar, 100 µm.

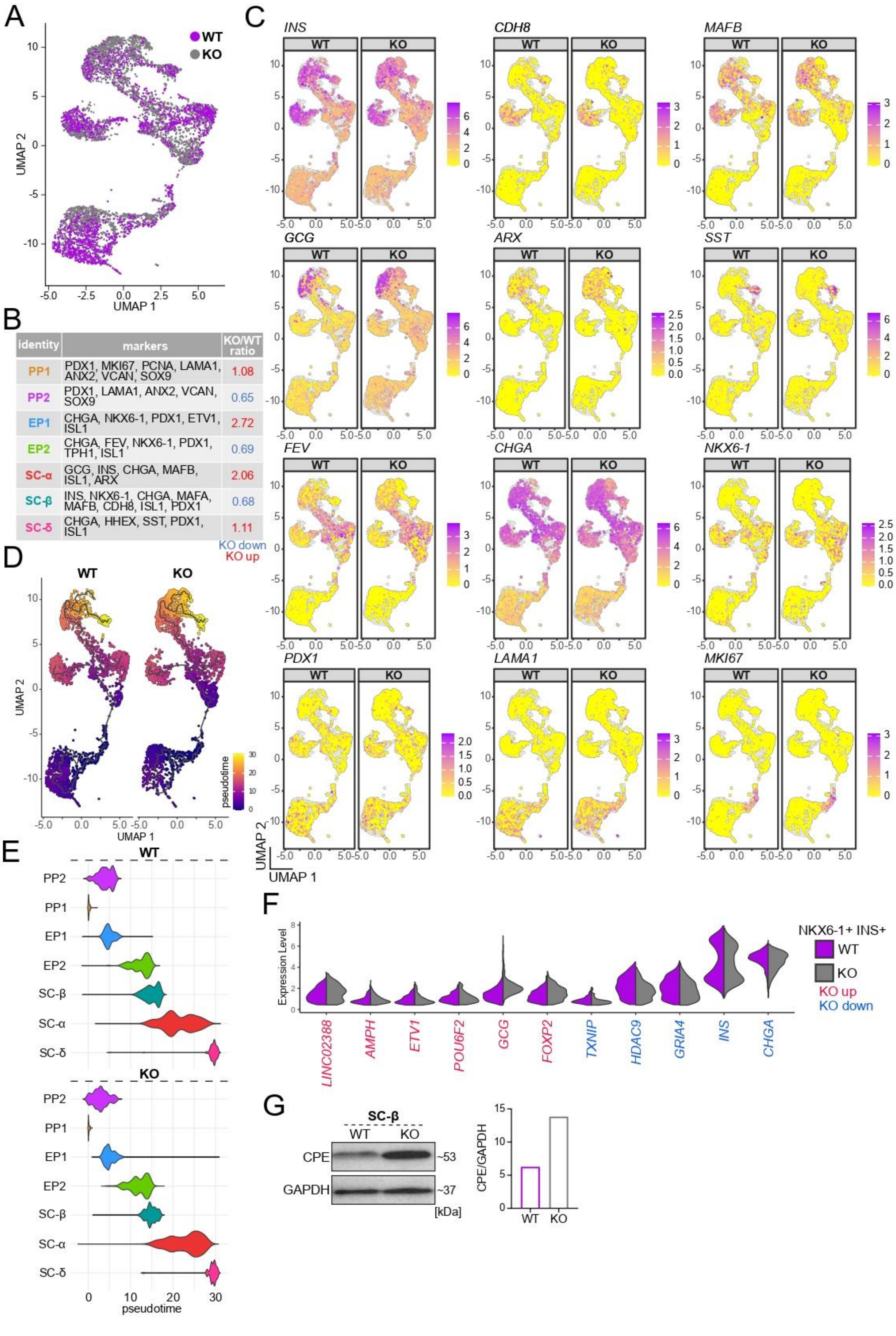
A
WT
KO
UMAP 1
UMAP 2
B
identity
markers
KO/WT ratio
PP1
PDX1, MKI67, PCNA, LAMA1, ANX2, VCAN, SOX9
1.08
PP2
PDX1, LAMA1, ANX2, VCAN, SOX9
0.65
EP1
CHGA, NKX6-1, PDX1, ETV1, ISL1
2.72
EP2
CHGA, FEV, NKX6-1, PDX1, TPH1, ISL1
0.69
SC-α
GCG, INS, CHGA, MAFB, ISL1, ARX
2.06
SC-β
INS, NKX6-1, CHGA, MAFA, MAFB, CDH8, ISL1, PDX1
0.68
SC-δ
CHGA, HHEX, SST, PDX1, ISL1
1.11
KO down
KO up
C
INS
CDH8
MAFB
GCG
ARX
SST
FEV
CHGA
NKX6-1
PDX1
LAMA1
MKI67
WT
KO
UMAP 1
UMAP 2
D
WT
KO
pseudotime
UMAP 1
UMAP 2
E
WT
PP2
PP1
EP1
EP2
SC-β
SC-α
SC-δ
KO
pseudotime
F
Expression Level
LINC02388
AMPH
ETV1
POU6F2
GCG
FOXP2
TXNIP
HDAC9
GRIA4
INS
CHGA
NKX6-1+ INS+
WT
KO
KO up
KO down
G
SC-β
WT
KO
CPE
GAPDH
~53
~37
[kDa]
CPE/GAPDH
WT
KO

**Supplementary Figure 2: Single-cell RNA-seq analysis of WT and FKBP2 cells at day 31 of pancreatic differentiation**

**(A)** UMAP plots of single-cell RNA sequencing (scRNA-seq) data from WT and FKBP2 KO cells at day 31 of differentiation.

**(B)** Table presents the key markers used to define each cellular cluster and the associated KO/WT ratio within each cluster.

**(C)** Feature plots displaying the mRNA expression of key markers: *INS*, *CDH8, MAFB*, *GCG*, *ARX*, *SST*, *FEV*, *CHGA*, *NKX6-1*, *PDX1*, *LAMA1*, and *MKi67*, across different pancreatic cell clusters for WT and KO samples at day 31 of differentiation.

**(D)** UMAP feature plot shows pseudotime values calculated using the Moncole3 package. WT and KO cells are ordered in pseudotime, starting from earlier cells, marked in blue, to later cells, marked in yellow. Each dot represents a single cell.

(**E**) Violin plot represents the pseudotime distribution of each cluster in WT and KO samples, positioned along the pseudotime trajectory.

(**F**) Violin plots comparing the mRNA expression of selected genes in WT and KO NKX6-1+/INS+ SC-β cells.

(**G**) Western blot analysis and quantification of carboxypeptidase E (CPE) protein levels in WT and KO SC-β cells. GAPDH served as a loading control. N = 2 independent biological replicates.

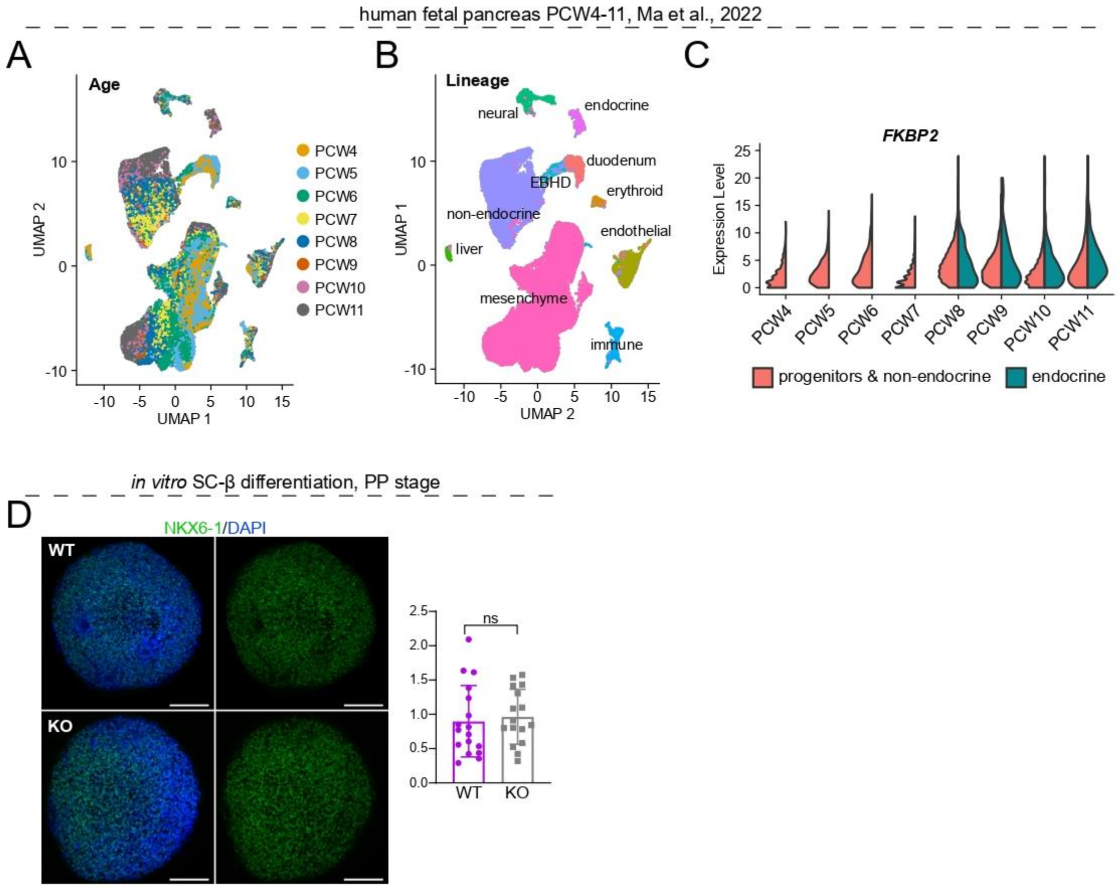


**Supplementary Figure 3: Single-cell RNA sequencing analysis of human fetal pancreas and immunofluorescence staining of FKBP2-deficient developing pancreatic SC-β cells**

**(A)** UMAP plot of human fetal pancreas cells (PCW4–PCW11 [42]). The plot shows the distribution of cells colored by post-conception week (PCW), ranging from week 4 (PCW4) to week 11 (PCW11).

**(B)** UMAP plot of the same dataset, with cells colored by their identified lineage. The major cell populations are labeled, including endocrine, neural, duodenum, erythroid, endothelial, non-endocrine, liver, mesenchyme, and immune cells.

**(C)** Violin plots showing the mRNA expression level of *FKBP2* across different developmental timepoints (PCW4–PCW11). The cells are categorized into two groups: "progenitors & non-endocrine" (red) and "endocrine" (teal). The plot demonstrates that FKBP2 expression is present in both cell types, with a trend of increasing expression in endocrine cells as gestation progresses.

**(D)** Immunofluorescence staining for NKX6-1 (green) in WT and FKBP2 KO at day 12, PP stage. DAPI marks in blue. Quantification (right) shows no significant difference (ns) in the NKX6-1+ area between WT and KO. The bar graph shows the quantification of the mean fluorescent intensity (MFI). Data are presented as mean ± SD, with each dot representing a single image. N = 3 independent experimental repeats, and an unpaired t-test was used for statistical analysis. Scale bar, 100 µm.

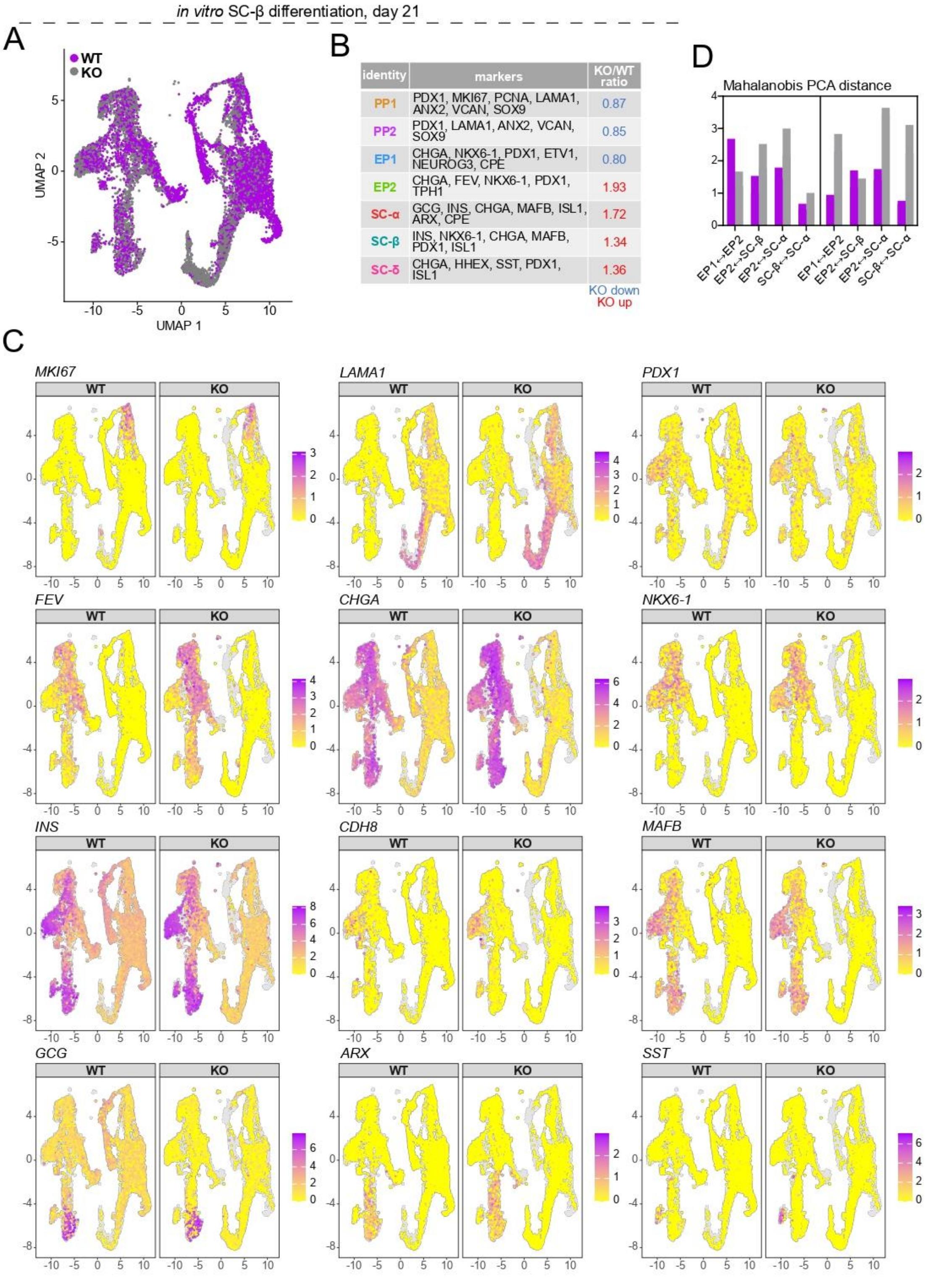
in vitro SC-β differentiation, day 21
A
WT
KO
UMAP 2
UMAP 1
B
identity
markers
KO/WT ratio
PP1
PDX1, MKI67, PCNA, LAMA1, ANX2, VCAN, SOX9
0.87
PP2
PDX1, LAMA1, ANX2, VCAN, SOX9
0.85
EP1
CHGA, NKX6-1, PDX1, ETV1, NEUROG3, CPE
0.80
EP2
CHGA, FEV, NKX6-1, PDX1, TPH1
1.93
SC-α
GCG, INS, CHGA, MAFB, ISL1, ARX, CPE
1.72
SC-β
INS, NKX6-1, CHGA, MAFB, PDX1, ISL1
1.34
SC-δ
CHGA, HHEX, SST, PDX1, ISL1
1.36
KO down
KO up
D
Mahalanobis PCA distance
EP1↔EP2
EP2↔SC-β
EP2↔SC-α
SC-β↔SC-α
C
MKI67
LAMA1
PDX1
FEV
CHGA
NKX6-1
INS
CDH8
MAFB
GCG
ARX
SST
WT
KO

**Supplementary Figure 4: Transcriptomic analysis of FKBP2 KO at day 21 of SC-β cell differentiation**

**(A)** UMAP plot of single-cell transcriptomes from SC-β cell differentiation (day 21) comparing WT (magenta) and FKBP2 KO (gray) cells.

**(B)** Cluster annotation identifies distinct cell populations: pancreatic progenitors (PP1, PP2), endocrine progenitors (EP1, EP2), β cells (SC-β), α cells (SC-α), and δ cells (SC-δ), along with representative marker genes and KO/WT abundance ratios. Clusters enriched in KO are highlighted in red, and those decreased in KO are highlighted in blue.

**(C)** UMAP feature plots showing mRNA expression of selected marker genes across WT and KO conditions, including proliferation (*MKI67*), basement membrane (*LAMA1*), pancreatic progenitor markers (*PDX1*, *NKX6-1*), endocrine and β cell markers (*CHGA*, *INS*, *MAFB,* and *CDH8*), endocrine progenitor marker (*FEV*), α cell markers (*ARX*, *GCG*), and δ cell marker (*SST).* The color intensity represents the expression level of each gene, from low (yellow) to high (purple).

**(D)** Bar graphs comparing Mahalanobis PCA distances between cell clusters at day 21 (D21, left panel) and day 31 (D31, right panel) for WT and KO cells. The distances reveal differences in the differentiation trajectories between WT and KO cells. For example, the distance between the EP2 and SC-α clusters is larger in KO cells, indicating altered progression towards the endocrine cell fate.

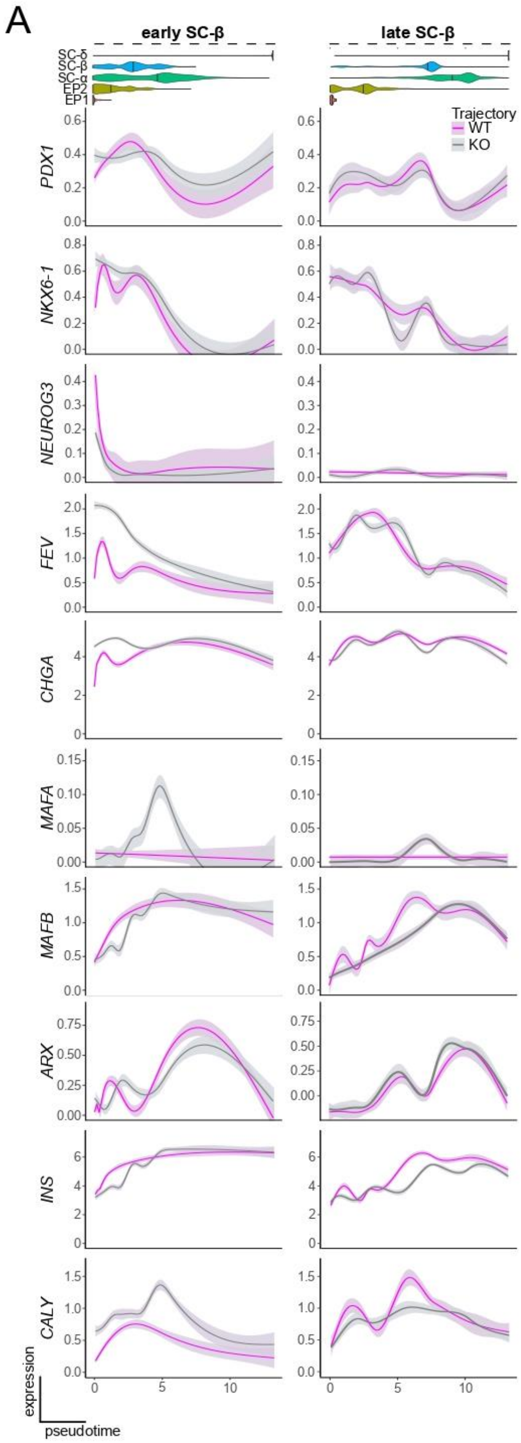


**Supplementary Figure 5: Expression dynamics of transcription factors and hormones along pseudotime during early and late SC-β cell differentiation**

**(A)** Expression dynamics along pseudotime trajectories for key transcription factors and hormones in both early and late SC-β cell differentiation. Plots show expression levels of *PDX1*, *NKX6-1*, *NEUROG3*, *FEV*, *CHGA*, *MAFA*, *MAFB*, *ARX*, *INS*, *CALY* across pseudotime for WT and KO conditions, with separate trajectories indicated for early and late SC-β cell populations.

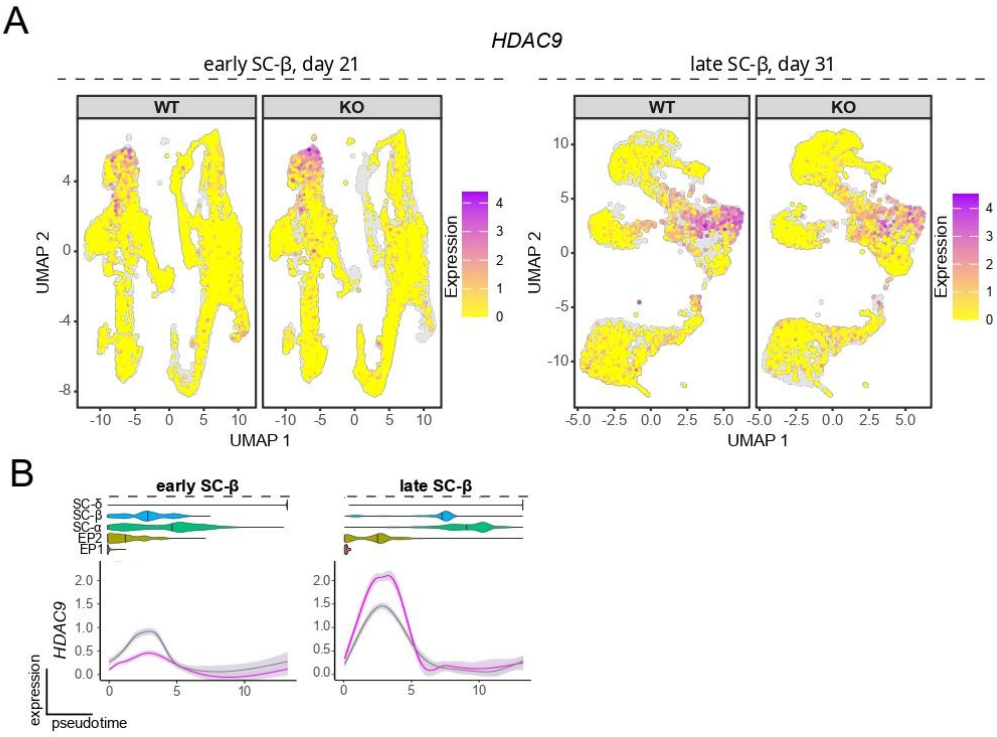


**Supplementary Figure 6: HDAC9 mRNA expression in early and late WT and KO SC-β cells**

**(A)** UMAP feature plots showing expression of *HDAC9* in WT and FKBP2 KO cells at two timepoints of SC-β differentiation: day 21 (early) and day 31 (late). Color intensity represents expression levels, from low (yellow) to high (purple).

**(B)** Expression dynamics along pseudotime trajectories for key transcription factors and hormones in both early and late SC-β cell differentiation. Plots show expression levels of *HDAC9* across pseudotime for WT and KO conditions, with separate trajectories indicated for early and late SC-β cell populations.

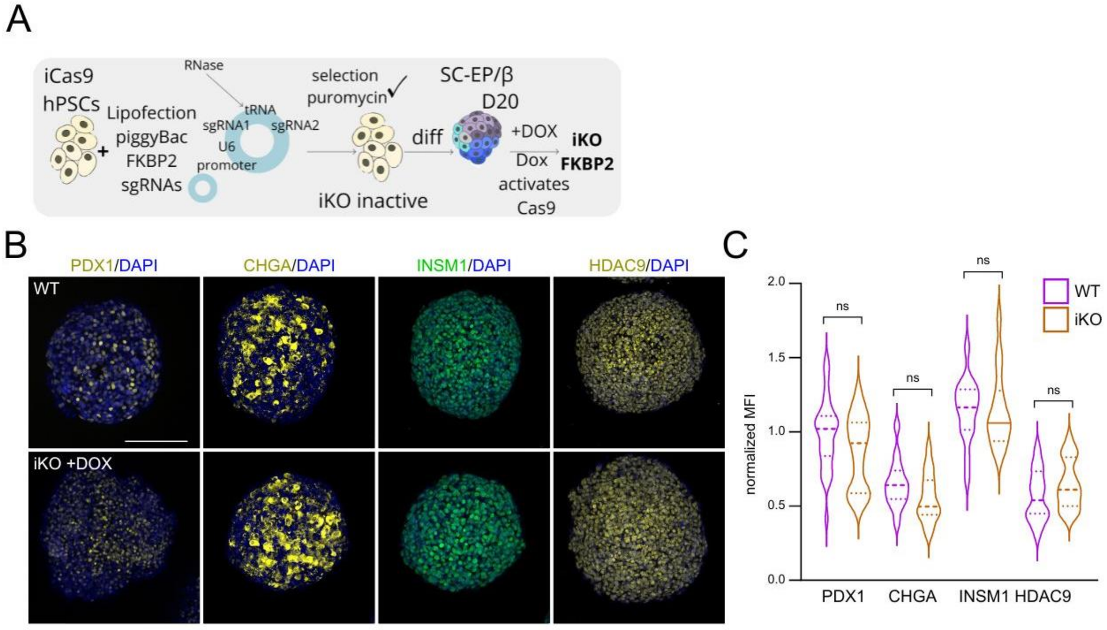


**Supplementary Figure 7: Inactivation of FKBP2 expression at SC-β cells does not affect β cell differentiation**

**(A)** Scheme of design to generate the doxycycline-inducible FKBP2 KO hPSCs. Human pluripotent stem cells (hPSCs) carrying an inducible Cas9 (iCas9) were transfected by lipofection with a PiggyBac vector encoding FKBP2-specific sgRNAs expressed under the U6 promoter. The construct contains tandem sgRNAs separated by a tRNA sequence to release individual sgRNAs. Following puromycin selection, stable hPSC clones harboring the inducible knockout cassette (iKO inactive) were obtained and subsequently differentiated toward EPs. On day 20 of differentiation, doxycycline (DOX) treatment activated Cas9 expression, resulting in sgRNA-guided cleavage and inducible knockout (iKO) of FKBP2 in SC-β cells.

**(B)** Immunofluorescence staining for PDX1 (yellow), CHGA (yellow), INSM1 (green), and HDAC9 (yellow) in WT and FKBP2 iKO at day 26, SC-β cell stage. DAPI marks nuclei in blue. Scale bar, 100 µm.

**(C)** Quantification of mean fluorescent intensity (MFI) for PDX1, CHGA, INSM1, and HDAC9 in WT and iKO cells, at day 26. N = 3 independent biological replicates. The violin displays the distribution of mean fluorescence intensity (MFI) normalized to DAPI, with the median (dashed line) and IQR (dotted line, representing 25th–75th percentiles). P-values were calculated using an unpaired Student t-test.

**Supplementary Table 1: Differentially expressed genes in KO vs WT SC-β cells at day 21 based on scRNA-seq.** The table includes information on: log2 fold change (log_fc), p-values (p_val, padj), expression levels in both groups (rate1, rate2), feature, and the compared groups (group1, group2).

**Supplementary Table 2: Differentially expressed genes in KO vs WT SC-β cells at day 31 based on scRNA-seq.** The table includes additional columns such as average expression (ave_expr) and test statistic (stat), in addition to log_fc, p-values (p_val), adjusted p-values (padj), expression levels in both groups (rate1, rate2), and feature. The comparisons include WT and KO samples at the late stage of differentiation.

**Supplementary Table 3: Table summarizes statistically significant gene expression changes between KO and WT SC-β cells at day 21 based on scRNA-seq.** It includes gene identifiers (feature), log2 fold change (log2FC), adjusted p-values (padj), average expression levels (rate.mean), cluster assignment (cluster), and regulation direction (direction).

**Supplementary Table 4: Table summarizes statistically significant gene expression changes between KO and WT SC-β cells at day 31 based on scRNA-seq.** It includes gene identifiers (feature), log2 fold change (log2FC), adjusted p-values (padj), average expression levels (rate.mean), cluster assignment (cluster), and regulation direction (direction).

**Supplementary Table 5:** Supplementary table to Fig. 2E, summarizing KEGG and GO pathway enrichment analysis of differentially expressed genes (DEGs) at day 31 in KO versus WT SC-β cells. Downregulated genes are mainly associated with ER, Golgi, secretory granules, and protein folding/secretion, while upregulated genes are linked to cell adhesion, cytoskeleton organization, calcium signaling, and cell-surface remodeling.
